# Popari: Modeling multisample variation in spatial transcriptomics

**DOI:** 10.1101/2025.05.08.652741

**Authors:** Shahul Alam, Tianming Zhou, Ellie Haber, Benjamin Chidester, Sophia Liu, Fei Chen, Jian Ma

**Affiliations:** Ray and Stephanie Lane Computational Biology Department, School of Computer Science, Carnegie Mellon University, Pittsburgh, PA 15213, USA; Machine Learning Department, School of Computer Science, Carnegie Mellon University, Pittsburgh, PA 15213, USA; Biophysics Program, Harvard University, Boston, MA 02115, USA; Harvard-MIT Division of Health Sciences and Technology, Massachusetts Institute of Technology, Cambridge, MA 02139, USA; Broad Institute of MIT and Harvard, Cambridge, MA 02142, USA; Ragon Institute of MGH, MIT and Harvard, Cambridge, MA 02139, USA; Department of Stem Cell and Regenerative Biology, Harvard University, Cambridge, MA 02138, USA

## Abstract

Integrating spatially-resolved transcriptomics (SRT) across biological samples is essential for understanding dynamic changes in tissue architecture and cell-cell interactions *in situ*. While tools exist for multisample single-cell RNA-seq, methods tailored to multisample SRT remain limited. Here, we introduce Popari, a probabilistic graphical model for factor-based decomposition of multisample SRT that captures condition-specific changes in spatial organization. Popari jointly learns spatial metagenes – linear gene expression programs – and their spatial affinities across samples. Its key innovations include a differential prior to regularize spatial accordance and spatial downsampling to enable multiresolution, hierarchical analysis. Simulations show Popari outperforms existing methods on multisample and multi-resolution spatial metrics. Applications to real datasets uncover spatial metagene dynamics, spatial accordance, and cell identities. In mouse brain (STARmap PLUS), Popari identifies spatial metagenes linked to AD; in thymus (Slide-TCR-seq), it captures increasing colocalization of V(D)J recombination and T cell proliferation; and in ovarian cancer (CosMx), it reveals sample-specific malignant-immune interactions. Overall, Popari provides a general, interpretable framework for analyzing variation in multisample SRT.

## Introduction

Recently, spatially-resolved transcriptomics (SRT) technologies – which jointly quantify and localize RNA transcripts via sequencing or imaging – have become essential tools for studying the *in situ* organization and interaction of cells across diverse contexts and biological processes [1, 2]. As these technologies advance, experimental designs increasingly encompass multiple SRT samples. For example, replicates from the same tissue enhance statistical power [3], while spatial atlases reveal transcriptomics diversity across developmental stages, anatomical regions, or disease states [4–11]. However, integrative analysis of multisample SRT (mSRT) presents new computational challenges, as methods designed for single slices are often inadequate in multisample settings [12–14]. In particular, there is a critical need for interpretable embedding methods that disentangle latent gene expression programs while capturing variation in their spatial patterns across samples. Such methods could link spatial transcriptomic heterogeneity to phenotypical variation, offering deeper insights into how spatial gene programs shape cellular function in health and disease.

Many computational tools exist for analyzing single-sample SRT. Methods such as MEFISTO [15], NSF [16], and SpiceMix [17] use factor-based models to infer spatial components of biological identity. MEFISTO and NSF incorporate spatial information via Gaussian processes, while SpiceMix leverages spatial proximity in a probabilistic graphical model. NSF and SpiceMix also enforce non-negativity, a key constraint for interpreting count-based SRT data. However, none are readily applicable to multisample contexts. For example, while MEFISTO models smoothness across spatial and sample coordinates, it cannot accommodate scenarios in which the differences between sample conditions are categorical.

Recent methods address specific facets of mSRT integration. PRECAST [18] uses a probabilistic factor model to correct batch effects. INSPIRE [19], STAligner [20], and spatiAlign [21] combine GNNs with contrastive learning to align sample embeddings. Other approaches – such as PASTE/PASTE2 [22, 23], CAST [24], and GPSA [25] – align samples to a common coordinate framework using optimal transport, regularized GNNs, or deep Gaussian processes. MOSCOT [26] and DeST-OT [27] extend optimal transport to model temporal dynamics. While these methods address alignment and batch correction, none provide a general, interpretable framework for analyzing spatial variation across samples. GNN-based methods like STAligner, spatiAlign, and CAST capture complex non-linearities that complicate interpretation. INSPIRE applies nonnegative matrix factorization (NMF) after a GNN encoder, but this post hoc design inherits the encoder’s non-linearities. Optimal transport-based methods enable alignment but lack gene- or factor-level attributions. PRECAST includes a factor-based model for batch correction but does not model spatial relationships between learned factors and lacks non-negativity constraints.

Here, we introduce Popari, a probabilistic, factor-based model that uncovers latent gene programs and quantifies their spatial affinities across samples in mSRT datasets. Unlike prior methods, Popari directly models variation in spatial affinities between gene programs and incorporates this information into both its likelihood and optimization. Applied to datasets profiling Alzheimer’s disease (STARmap PLUS), thymic involution (Slide-TCR-seq), and ovarian cancer (CosMx), Popari uniquely captures sample-specific differences in metagene interactions and cell-cell accordance, offering a biologically interpretable lens on mSRT dynamics.

## Results

### Overview of Popari

Popari performs joint analysis of SRT samples from multiple conditions, timepoints, or individuals (**Fig.** 1A and **Methods**). It represents “spatial entities” – either biological cells (imaging-based SRT) or spatially-barcoded spots (sequencing-based SRT) – by embedding their gene expression profiles into an interpretable, low-dimensional latent space. Popari also learns embeddings at multiple spatial resolutions through spatial downsampling, enabling it to capture both fine-grained (e.g., cell type level) and coarse-grained (e.g., spatial domain level) variation in mSRT dataset. The model employs a probabilistic graphical model framework, where nodes correspond to spatial entities and edges encode spatial proximity, allowing it to detect shared and differential structures across resolutions and samples.

**Figure 1:**
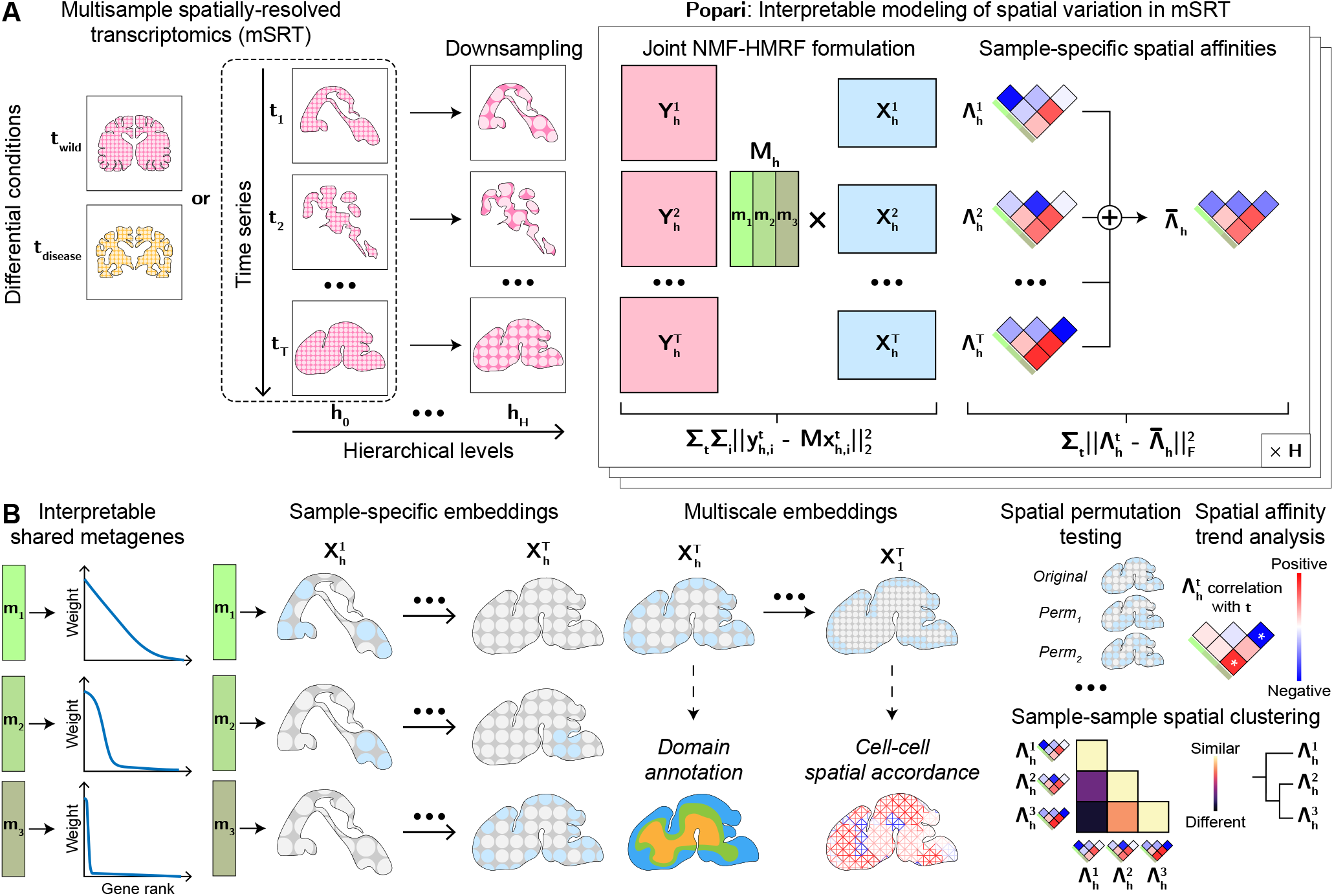
Overview of the Popari model. **A**. Popari takes as input *T* spatial transcriptomics samples with a temporal or other differential structure. For each sample *t* ∈ [1, …, *T*], the gene expression *Y* ^*t*^ is decomposed into two non-negative factors: shared metagenes *M* and latent embeddings *X*^*t*^. Spatial coordinates are used to define an *H*-level hierarchical neighborhood graph, with low-resolution nodes obtained via spatial downsampling. For every edge at the lowest level of the graph, spatial accordance between the corresponding nodes is modeled as the inner product of their embeddings, weighted by a sample-specific metagene affinity matrix Λ^*t*^. **B**. The learned *M, X*^*t*^, and Λ^*t*^ offer an interpretable, multiscale view of spatial transcriptomic variation. Down-stream analyses include spatial domain annotation, scoring of spatial accordance between single cells, spatial permutation testing, spatiotemporal dynamics analysis, and clustering of SRT samples by spatial structure.

At each spatial resolution *h*, the model infers *K* metagenes – shared fundamental expression factors across all *T* samples – represented by a *G* × *K* metagene matrix *M*_*h*_, where *G* is the number of genes. Each entity is embedded into the shared embedding space defined by *M*_*h*_; for entity *i* in sample *t* and resolution *h*, this embedding is 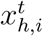, and the observed gene expression 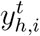 is modeled as 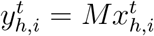, a linear, non-negative model. Crucially, for each sample, Popari also estimates a *K* × *K* spatial affinity matrix 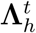 between metagenes, where each entry quantifies co-expression of metagene pairs in neigh-boring entities. Comparing 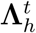 across samples reveals sample-specific differences in spatial metagenes’ structure.

Popari includes two user-defined hyperparameters: one controlling the influence of spatial information, and the other controlling expected sample-to-sample variation in 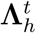. The metagenes *M*_*h*_, latent embeddings 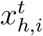, and spatial affinity matrices 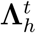 are jointly optimized via an alternating maximum a posteriori (MAP) optimization procedure. After optimization, Popari provides multiscale insights into spatial structure (**Fig.** 1B). Each column *m*_*k*_ of the metagene matrix represents a gene co-expression module with high-weight genes forming the core of the module. This offers a conceptual advantage over methods that detect individual spatially variable genes (SVGs), as metagenes capture nuanced, combinatorial gene expression patterns [17, 28]. Trends in 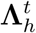 across samples highlight shifts in spatial coherence of metagene pairs. Permutation of the 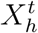 enables statistical significance testing of the inferred 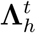 values. Additionally, given categorical annotations (e.g., cell types), Popari uses 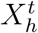 and 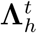 to compute “accordance scores” between each pair of categories. Here, “accordance” refers to the degree to which the metagene spatial affinities 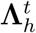 learned by the model agree with the correlation patterns of the learned 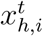 along edges of the spatial graph.

Further mathematical details, optimization procedures, implementation, and interpretation guidelines are provided in **Methods** and **Supplemental Information**. The Popari model, an efficient optimization framework, and a built-in hyperparameter grid search tool are available as a Python package on PyPI.

### Simulations demonstrate modeling advantages of Popari

To demonstrate Popari’s conceptual advantages, we evaluated it on simulated SRT data, identifying three essential criteria for the simulation framework: (1) a metagene-based generative model for gene expression counts; (2) a spatial mechanism for introducing dropout-based sparsity; and (3) an interface to define (m)SRT, allowing for sample-specific ground truth metagenes and spatial accordances. Existing methods for simulating SRT do not meet these criteria [16, 29–31], so we adapted a prior framework [17] by incorporating two novel components: (1) a hierarchical module, which samples independent sets from the spatial graph and drops out expression counts per gene; and (2) a differential module, which enables sample-specific metagene spatial distributions (**Fig.** S1). Although the modules can be used together, we design two separate simulations that each use only one, allowing us to isolate and evaluate Popari’s contributions. Additional details of the simulation modules are provided in the **Supplemental Information**.

We introduce two metrics to evaluate recovery of ground truth metagene spatial patterns. The *spatial Wasserstein distance* (SWD) measures the “spatial congruence” between ground truth and learned metagene embeddings by solving a minimum-cost flow problem, detecting incongruence where non-spatial metrics cannot (**Fig.** S2). The *affinity Spearman correlation* (ASC) matches metagenes using SWD and quantifies consistency of the inferred **Λ**^*t*^ parameters. We found these metrics to be more sensitive to spatial discrepancies than non-spatial metrics such as adjusted Rand index (ARI); see **Supplemental Information** for formal definitions.

We first evaluated Popari’s ability to overcome structured sparsity using the hierarchical module to generate an *in silico* brain cortex SRT dataset. For each simulated gene, 15% of entities had their expression of that gene set to zero. As an ablation, we introduce *hierarchical NMF*(NMF-H), which uses Popari’s spatial downsampling component but does not use spatial information during inference. Compared to NMF, SpiceMix, NSF, and NMF-H, Popari consistently recovered ground truth spatial metagenes under spatial dropout (**Fig.** 2A, top; **Fig.** 2B, left; and **Fig.** 2C). For example, Popari accurately recovered the spatial pattern of metagene 7 (corresponding to the L2 layer; **Fig.** 2C), whereas SpiceMix failed to identify metagenes with clear spatial structure. NMF and NMF-H identified all metagenes but did not utilize spatial information for denoising. NSF produced oversmoothed embeddings that failed to capture the simulated layer-like structures. Quantitatively, Popari achieved a lower average SWD (0.12) of all other methods (**Fig.** 2A, top; NMF: 0.29; NMF-H: 0.24; SpiceMix: 0.20; and NSF: 0.31). It also achieved a higher ASC (0.72) than SpiceMix (0.31), confirming better recovery of ground truth **Λ**^*t*^ values (**Fig.** 2B, left). Popari’s advantage in mSWD also held at coarser hierarchical levels (**Fig.** S3A).

**Figure 2:**
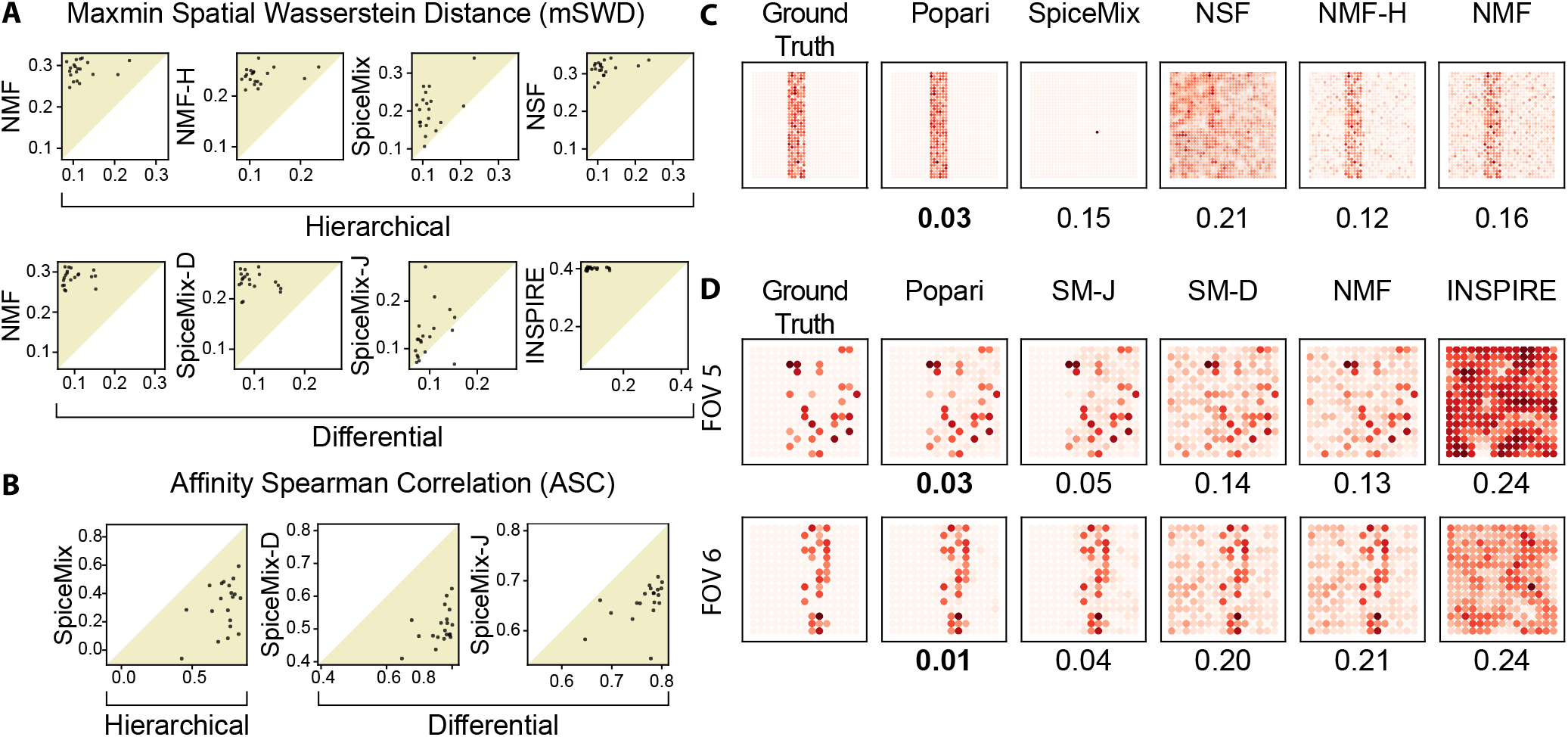
Benchmarking on simulations of (m)SRT. **A**. Performance comparison between Popari and baseline methods using the max-min spatial Wasserstein distance (mSWD) metric (see **Supplemental Information**). Each dot represents paired scores from 20 simulation replicates (Popari: *x* -axis); baseline: *y* -axis), with scores selected from the best of five random initializations. Tan-colored half-planes indicate regions where Popari outperforms the baseline. **B**. Similar performance comparison as in **(A)**, using the affinity Spearman correlation (ASC) metric. **C**. *In situ* embeddings of metagene 7 at hierarchical level *h* = 0 in the hierarchical simulation. SWD scores between the embedding of each method and ground truth are shown below each square. **D**. *In situ* embeddings of metagene 9 from the differential simulation for two FOVs (FOV 5, top; FOV 6, bottom), with corresponding SWD values shown below.

Next, we evaluated Popari’s performance in modeling multisample spatial variation using the differential module to generate ten simulated brain cortex SRT samples. The samples, split into two groups, shared cell types (modeled after real glial, excitatory, and inhibitory subtypes) and spatial domains (modeled after cortical layers) but differed in cell type proportions across spatial domains, leading to variation in spatial accordances of ground truth metagenes. We compared Popari to NMF, INSPIRE, and two SpiceMix variants: *disjoint* SpiceMix (SM-D), which learns separate **Λ**^*t*^ for each sample without differential regularization; and *joint* SpiceMix (SM-J), which uses a shared **Λ** across samples. Popari outperformed all methods in recovering spatial metagene embeddings (**Fig.** 2A, bottom, **Fig.** 2B, right, and **Fig.** 2D). For example, Popari accurately recovered metagene 9, associated with a layer-specific excitatory cell type in one group and a diffuse pattern in the other, whereas other methods failed to denoise its spatial embedding pattern **Fig.** 2D). Popari achieved the lower average SWD (0.10) than other methods (**Fig.** 2A, bottom; NMF: 0.29; INSPIRE: 0.40; SpiceMix-D: 0.24; and SpiceMix-J: 0.13) and the highes ASC (0.76) compared to SM-D (0.51) and SM-J (0.65), confirming improved recovery of sample-specific spatial affinities (**Fig.** 2B, right). Visual comparisons of ground truth, learned, and empirical affinities further illustrate Popari’s advantage (**Fig.** S3B).

Overall, supported by our simulation framework, these results demonstrate the conceptual advantage of Popari in recovering spatial structure in SRT under both sparsity and multisample variation.

### Revealing spatially variable components of Alzheimer’s Disease

We applied Popari to an mSRT dataset profiling eight coronal brain sections from TauPS2APP mouse models of Alzheimer’s disease (AD) and age-matched controls at 8 and 13 months, generated using the STARmap PLUS platform. The dataset includes single-cell expression measurements for 2,766 genes across 72,165 cells [32], along with *in situ* spatial localization of amyloid beta (A*β*) plaques and hydrophoshorylated tau (p-tau) tangles, which are key proteins implicated in AD pathologies [33, 34]. Popari recapitulates known neuronal and glial cell types while capturing AD-associated changes in their spatial organization.

Popari’s embedding space clearly separates cell types identified in the original study (**Fig.** S4). Glial subtypes – astrocytes (Astro), microglia (Micro), and oligodendrocytes (Oligo) – are particularly well-separated, a critical distinction given their AD-specific subtypes [35–37]. Clustering the embedding space reveals consistent spatial domains across replicates (**Fig.** S5A), with marker gene analysis aligning these domains to known brain regions (**Fig.** S5B). Mapping metagenes to cell types based on average expression (**Fig.** 3A) shows, for example, that metagene m2 is strongly expressed in glial cells, while metagenes m7, m4, and m9 are specific to Micro, Oligo, and Astro cells, respectively.

**Figure 3:**
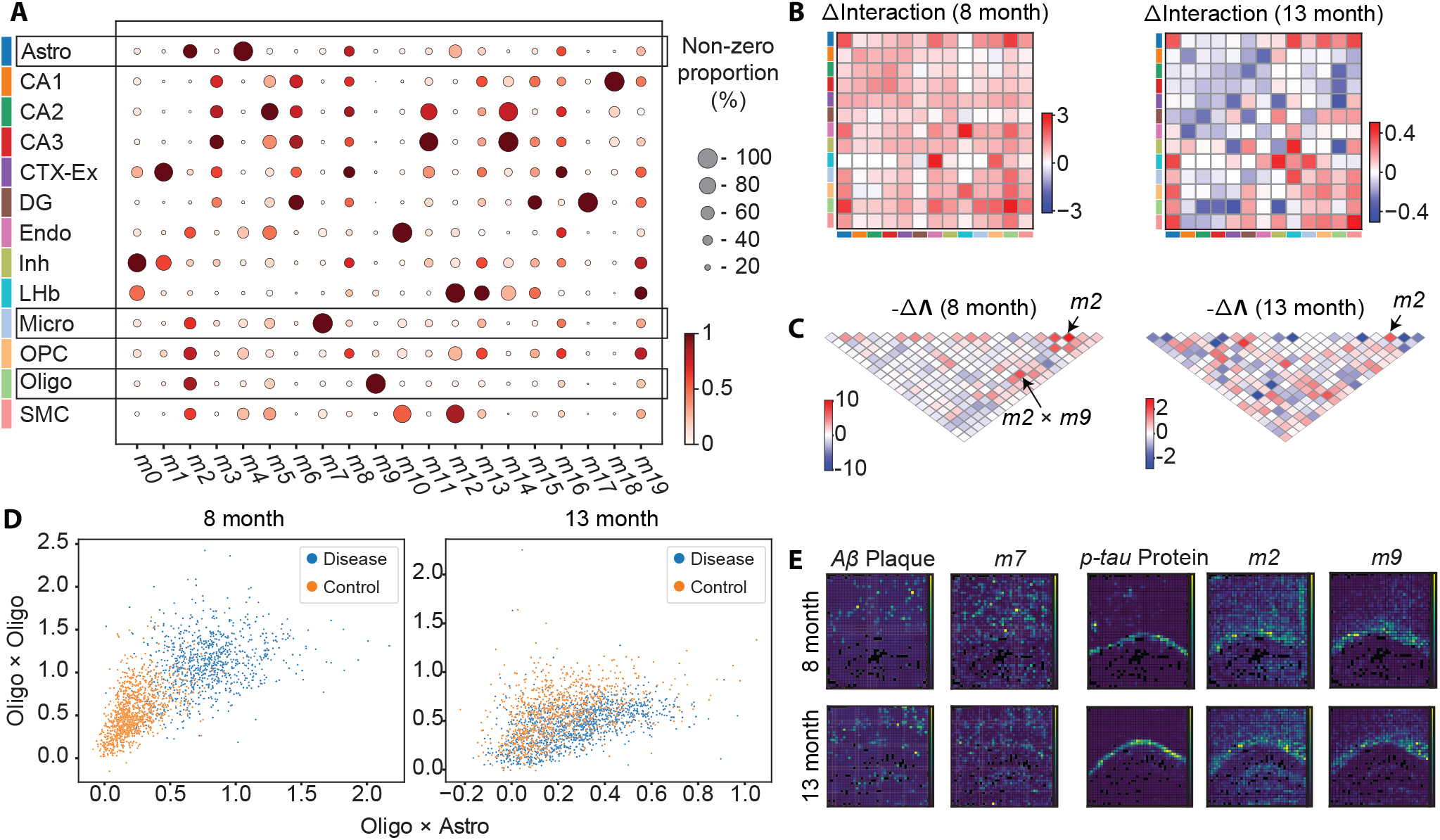
Revealing spatially variable components of Alzheimer’s disease from STARmap PLUS data. **A**. Correspondence between cell types and average metagene expression, with glial cell types boxed. **B**. Δ*U*_*x*_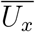 score matrices showing changes in cell-type accordance strength between control and disease samples at 8 months (left) and 13 months (right). Cell type color key matches **A. C**. -Δ**Λ** matrices comparing disease and control samples at 8 months (left) and 13 months (right). **D**. 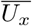 scores for individual oligodendrocytes with respect to Oligo–Astro accordances (x-axis) and Oligo–Oligo self-accordances (y-axis) at 8 months (left) and 13 months (right). **E**. *In situ* expression of A*β* plaque (first column) and *p-tau* protein (third column), juxtaposed with *in situ* learned embedding values of metagenes m7, m2, and m9 (second, fourth and fifth columns) for 8 months (top row) and 13 months (bottom row). Values shown are for entities at hierarchical level *h* = 1.

To assess disease-related changes in spatial organization, we compared cell-type accordance across TauPS2APP and control samples. At 8 months, Astro-Astro, Oligo-Oligo, and Oligo-Astro pairs show the largest increase in accordance scores 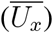(**Fig.** 3B). 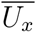 quantifies the average value of the edge energy function *U*_*x*_ for edges between those cell types; see **Supplemental Information** for details. By contrast, 13-month samples show smaller overall changes, with self-accordance of cortex excitatory (CTX-EX) and dentate gyrus (DG) neurons decreasing the most. Visualization of individual cell 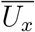 scores confirms that oligodendrocytes in 8-month diseased samples exhibited elevated Oligo-Oligo and Oligo-Astro accordances, a pattern less distinct at 13 months (**Fig.** 3C). Spatial plots further localize these changes: elevated Oligo-Oligo and Oligo-Astro accordances appear in the white matter at 8 months, while Oligo-CTX-EX accordances are elevated in the neocortex at 8 months but strongly reduced by 13 months (**Fig.** S5C). We next examined differences in metagene spatial affinities between TauPS2APP and control samples, quantified by Δ**Λ** (**Fig.** 3D). At 8 months, metagene m2 shows the largest increase in self-affinity, and both m2 and m9 exhibit stronger cross-affinities. These changes are less pronounced at 13 months, although m2’s self-affinity remains elevated in diseased samples compared to controls.

To further validate metagene-disease associations, we examined spatial correlations between metagenes and AD pathology markers. Metagenes m2 and m9 show the strongest correlation with p-tau, while m7 correlates with A*β* plaque (**Fig.** 3E). Signature genes of these metagenes overlap with known AD-associated gene modules (**Fig.** S6). The m2 signature significantly overlaps with disease-associated astrocyte (DAA; 44.44% overlap, Fisher’s exact *P* = 4.56 × 10^*−*2^) and disease-associated microglia (DAM; 37.04% overlap, *P* = 7.91 × 10^*−*6^) gene modules. The m7 signature overlaps with plaque-induced gene (PIG) modules from the original study, including established PIGs such as *Apoe, Ctsb, C1qb, Cd63*, and *Gfap* (*P* = 1.29 × 10^*−*4^ at 8 months; *P* = 1.69 × 10^*−*4^ at 13 months) [38]. All genes overlapping between m7 and the 8-month PIG set also appear in the 13-month set, suggesting that m7 captures persistent plaque-induced gene expression in later disease stages.

Thus, Popari simultaneously recapitulates high-resolution cell types and low-resolution spatial expression patterns in STARmap PLUS data while inferring differential metagene spatial organization between the AD and control conditions. These results highlight Popari’s capacity for comparative analysis of spatial structure between phenotypic states.

### Characterizing temporal variation in thymic architecture and T cell organization

We next applied Popari to explore mouse thymic involution, the process of gradual thymus shrinkage during aging, which is correlated with decreased production of naïve T lymphocytes and reduced immune system robustness [39]. This evolutionarily conserved process occurs across nearly all vertebrates, making a spatially-resolved understanding of involution key to uncovering the drivers of age-related immune decline. Popari was applied to a five-sample Slide-TCR-seq dataset from mouse thymuses at 3, 28, 35, 91 and 105 days post-conception [40], and effectively identified spatiotemporal changes in thymic organization at the level of metagenes, cell types, and domains.

Using low-resolution bin embeddings, Popari discerned key spatial domains across all timepoints, clearly distinguishing the cortex and medulla, the two primary thymic regions (**Fig.** S7A). Popari also identified concentrically arranged sub-domains within both the medulla and cortex, consistent with the “continuous cortico-medullary axis” described in prior work [41]. *In situ* visualization of metagene embeddings showed that some metagenes are restricted to either cortex or medulla, while others exhibit more complex spatial patterning (**Fig.** S8). We further examined the composition of the learned metagenes *M* by annotating each Slide-TCR-seq bead using robust cell type decomposition (RCTD) [42] and an annotated single-cell RNA-seq thymus atlas [43]. Average metagene expression per cell type (**Fig.** S7B) revealed distinct mapping – for example m5 maps to medullary thymic epithelial cells (mTECs), m10 to cortical thymic epithelial cells (cTECs), and m17 to effector lymphocytes (ELCs). Enrichment plots showed that m17 is particularly enriched in regulatory T cells (Treg), while m4 is enriched in proliferating double-positive (DP(P)) and, secondarily, double-negative (DN(P)) T cells (**Fig.** S9). The mapping between metagenes and classes is not one-to-one: for example, m1, m4 and m11 are primarily expressed in thymocytes (TCs) and secondarily in cTECs, while m12 is primarily expressed in antigen-presenting cells (APCs) and secondarily in mTECs.

Popari also captured temporal changes in spatial accordances between metagene pairs through timepoint-specific affinity parameters **Λ**^*t*^, measured by Pearson correlation of pairwise affinity with mouse age (**Fig.** 4A). Metagenes m6 and m8 exhibited strong positive temporal correlations in self-affinity (r=0.81 with slope=42.96; and r=0.54 with slope=41.87, respectively). Gene Ontology (GO) analysis [44] of top-ranked signature genes in each metagene revealed that m6 is linked to T cell differentiation processes (e.g., *MAPK* cascade regulation), while m8 is associated with immune response functions (e.g., NF-*κ*B signaling and viral entry inhibition) (**Fig.** S10). The strongest positive cross-affinity correlation was between m1 and m4 (r=0.72, slope=45.09); m1’s top genes are linked to V(D)J recombination, while m4’s genes are linked to T cell proliferation (e.g., *Tcf7* [45], *Lat* [46–48]). GO terms associated with m4 – including cell division, cell cycle, and mitotic chromatid separation – further support its role in proliferation. These findings suggest that V(D)J recombination and T cell proliferation become increasingly spatially coordinated over time. Conversely, m4 and m17 show the strongest negative correlation over time, indicating growing spatial anti-correlation between proliferating thymocytes and Tregs. At the cell-cell spatial accordance level, *U*_*x*_ scores varied over time (**Fig.** 4B). Most self- and cross-accordances between broad cell type categories moderately decreased, with the exception of stromal cells, which showed increased self-affinity. Fine-grained analysis revealed that this increase was primarily driven by plasmocytoid dendritic cells (pDCs), monocytes (Mono), and fibroblasts (Fb) **Fig.** S11).

**Figure 4:**
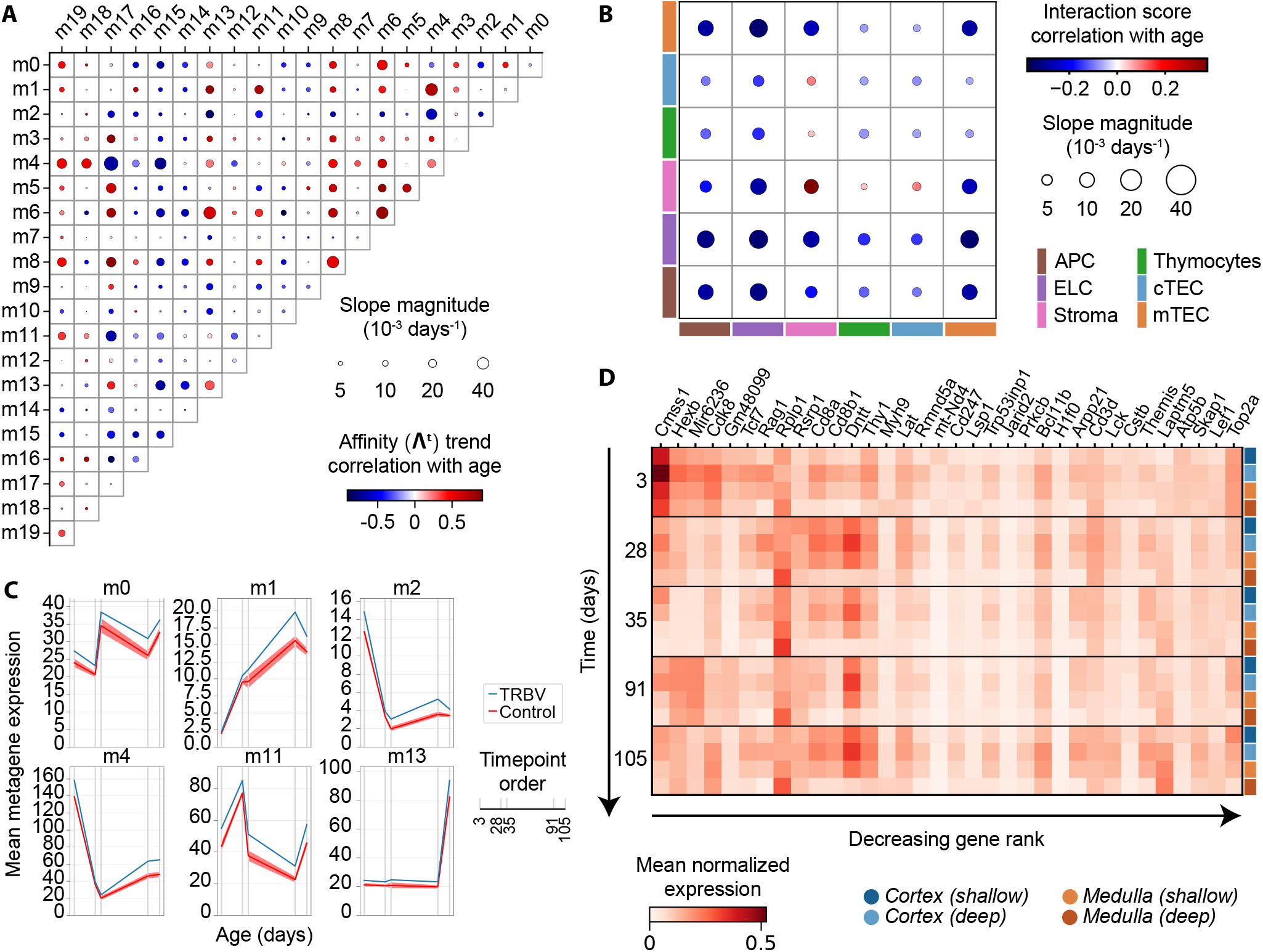
Characterizing temporal variation in thymic architecture and T cell organization. **A**. Pearson correlation and regression slope of spatial affinity values from the **Λ**^*t*^ matrices with age, at hierarchical level *h* = 1. **B**. Pearson correlation and regression slope of spatial accordance scores between broad cell type pairs with age, also at hierarchical level *h* = 1. **C**. Line plot showing mean embedding values in spots with non-zero T cell receptor beta-chain variable (TRBV) usage for selected metagenes. **D**. Average expression of top genes for selected metagenes (from **C**) across timepoints and spatial domains. Every group of four rows in the heatmap corresponds to a single timepoint (age increasing top to bottom); within each group, rows represent spatial domain ordered by depth along the corticomedullary axis. Cell type abbreviations: APC: antigen-presenting cells; ELC: effector lymphocytes; TC: thymocytes; cTEC: cortical thymic epithelial cells; mTEC: medullary thymic epithelial cells.

We further examined correlation between T cell receptor beta chain variable (TRBV) and joint (TRBJ) region usage (coassayed by Slide-TCR-seq) and Popari metagene expression. A subset of metagenes showed enriched expression in bins expressing TRBV regions across all timepoints (**Fig.** 4C), with similar trends for TRBJ-using (**Fig.** S12). These metagenes contained signature genes associated with V(D)J recombination and T cell development. Importantly, TRB usage was not included as an input to Popari, further validating its ability to uncover biological patterns across samples. We also assessed transcriptomic correlates of TRBV usage by plotting average expression of key genes across timepoints and spatial domains (**Fig.** 4D). Many of the V(D)J recombination-associated genes such as *Rag1, Top2a*, and *Dntt* exhibited spatial gradients consistent across timepoints [43, 49, 50]. An exception was *Cmss1*, a cell cycle gene with higher expression at day 3 than at later timepoints.

Overall, our analysis of Slide-TCR-seq thymus data with Popari successfully recovered known immune cell types and spatial domains while quantifying gradual architectural changes during aging, demonstrating the advantages of Popari’s multiscale, multisample modeling framework.

### Discerning malignant-immune spatial interaction in ovarian cancer

We further applied Popari to explore spatial heterogeneity across tumor samples from patients with tubo-ovarian high-grade serous carcinoma (HGSC). In contrast to prior settings, tumors exhibit marked diversity in tissue morphology and anatomical organization. This spatial complexity obscures understanding of the molecular determinants of cancer severity and immunotherapy responsiveness in ovarian cancer. To address this gap, we applied Popari to a CosMx spatial transcriptomics dataset consisting of 27 adnexal tumor samples comprising 161,962 cells, thereby accomplishing a differential analysis of spatially-resolved HGSC transcriptomes [51]. Popari successfully identified tumor-infiltration-associated metagenes, stratified tumor samples by degree of stromal/malignant compartmentalization, and summarized malignant cell state heterogeneity across the cohort.

Using hallmark gene programs from [51], we confirmed that shared Popari metagenes corresponded to known HGSC cell states. Notably, over half of the learned metagenes are enriched for malignant cell marker genes, corroborating inter-patient heterogeneity of malignant cell transcriptomes (**Fig.** 5A). On the other hand, m14 and m16 are associated with immune cell types such as T cells, NK cells, and monocytes. In particular, m14 is enriched for the tumor-infiltrating lymphocyte (TIL) program, which is exclusively expressed by lymphocytes proximal to malignant cells (**Fig.** S13A). Complementarily, metagenes m8, m14, and m2 are enriched for the TIL-adjacent malignant cell program (M_TIL_ (UP)), while m1, m6, and m9 are enriched for the non-TIL-adjacent malignant cell program (M_TIL_ (DOWN)). Metagene m0 is enriched for the fibroblast program, and m15 represents a program associated with fibroblast desmoplasia (**Fig.** S13B). Visualization of the Popari embeddings illustrates transcriptomic diversity across patients while also capturing shared non-malignant cell types (**Fig.** 5B).

**Figure 5:**
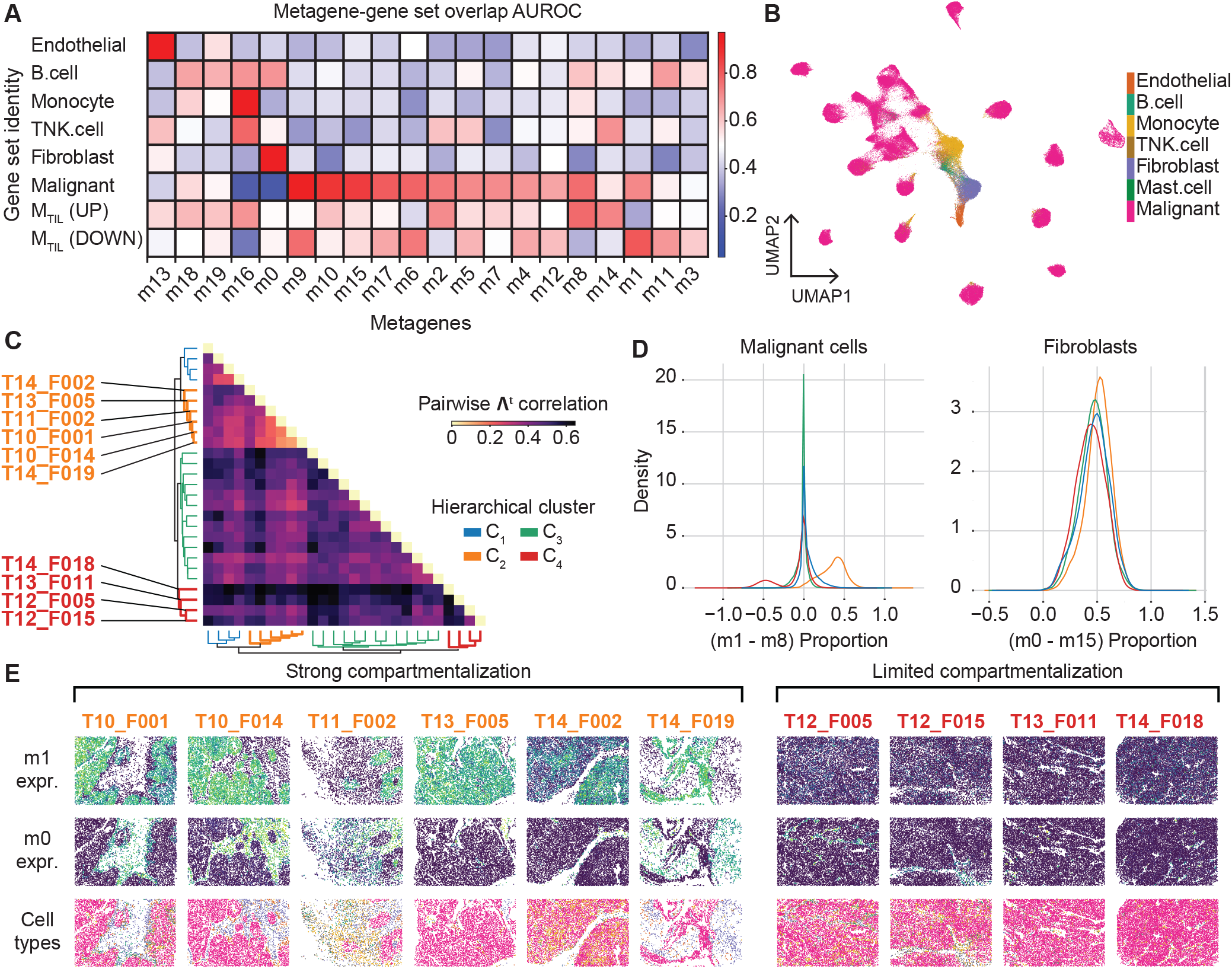
Discerning sample-specific characteristics of malignant-immune cell spatial interaction in ovarian cancer. **A**. AUROC scores capturing the alignment of learned Popari metagenes with predefined marker gene sets identified in [51]. **B**. UMAP visualization of Popari embeddings colored by cell types identified in [51]. **C**. Pairwise correlation between **Λ**^*t*^ matrices for all samples, with rows and columns sorted via hierarchical Ward clustering. The dendrogram at the bottom indicates the hierarchical relationship between samples. **D**. Kernel density estimate (KDE) for the distribution of metagene proportion differences for selected cell types, grouped by hierarchical cluster; malignant cells are plotted against the proportional difference score (m8 - m12) (left), while fibroblasts are plotted against the proportional difference (m0 - m15) (right). Cluster colors match those given in **C. E**. *In situ* expression of metagenes m0 and m1, alongside cell type labels, for samples in clusters *C*_2_ and *C*_4_ (identified in **C**.).

Beyond cell state characterization, Popari captured differences in spatial organization across samples. By clustering the learned spatial affinity matrices **Λ**^*t*^, we grouped tumor samples into four clusters (**Fig.** 5C). Clusters *C*_2_ and *C*_4_ exhibited the most distinct spatial structures. Samples in *C*_2_ showed strong self-affinities of both m0 (fibroblasts) and m1 (malignant cells), while this was absent in *C*_4_ (**Fig.** S14A). We examined these clusters further by comparing relative metagene proportions (see **Methods** for details). Malignant cells in *C*_2_ showed a greater difference between m1 (M_TIL_ (DOWN)) and m8 (M_TIL_ (UP)) proportions than those in *C*_4_ (**Fig.** 5D, left). Similarly, fibroblasts in *C*_2_ showed a greater difference between m0 (wild-type fibroblasts) and m15 (desmoplastic fibroblasts) than those in *C*_4_ (**Fig.** 5D, right). These patterns were further supported by differences in expression of top-weighted genes from these metagenes, which reflected the spatial affinity and abundance trends (**Fig.** S15). Thus, Popari distinguishes tumor samples by both spatial arrangement and relative abundance of fibroblast and malignant cell subtypes.

*In situ* visualization of the learned embeddings for metagenes m0 and m1 confirmed qualitative differences in their spatial distributions between clusters *C*_2_ and *C*_4_ (**Fig.** 5E, top and middle rows). When overlaid with *in situ* arrangement of cell types, m0 and m1 clearly delineate the stromal and malignant compartments, respectively (**Fig.** 5E, bottom row). Distinguishing this microenvironmental structure is critical for characterizing HGSC, as it is well-known that lymphocytes preferentially localize to the stromal compartment [51, 52]. The distinction between compartments is further supported by the *U*_*x*_pair score between m0 and m1 (see **Supplemental Information**); *in situ* visualization of this score along edges of the spatial graph highlights the compartment boundary, which is clearly defined in *C*_2_ samples but absent in *C*_4_ (**Fig.** S14B).

Overall, Popari successfully identifies shared malignant and non-malignant cell states in CosMx HGSC data, despite high intersample heterogeneity. It also distinguishes tumor samples based on differences in tumor compartmentalization and spatial organization, demonstrating Popari’s utility for comparative spatial analysis even in contexts with high biological variation between samples.

## Discussion

We developed Popari, an interpretable probabilistic model for uncovering the dynamics of spatially-aware metagenes from mSRT datasets. Popari’s core methodological contributions are: (1) a regularized model that captures differential spatial structures across samples; and (2) an integrative, hierarchical framework for modeling SRT at multiple spatial resolutions. Across simulated (m)SRT datasets, we introduced new spatial evaluation metrics and demonstrated that Popari consistently outperforms baseline methods in recovering ground truth spatial structure. On the imaging-based STARmap PLUS data, Popari delineated glial and neuronal cell types, identified AD-associated metagenes *de novo*, and revealed changes in spatial accordances between disease and control conditions. On Slide-TCR-seq data, Popari distinguished thymocytes and stromal cell types, detected temporally correlated shifts in spatial affinity among immune-related metagenes, and recovered spatial domain structure of the cortex and medulla. On CosMx tumor samples, Popari identified diverse malignant and immune cell states, stratified tumors by spatial organization, and resolved differences in the degree of tumor compartmentalization. These findings underscore the utility of Popari for spatial factor analysis in complex, heterogeneous tissue environments. By leveraging shared metagene structure while allowing sample-specific variation in spatial relationships, Popari is uniquely suited to uncover both conserved and condition-specific patterns in spatial omics data.

Despite its strengths, Popari has limitations and opens up future directions for development. One challenge is the presence of batch effects across samples. Although batch correction is a widely-explored issue in single-cell and spatial omics analysis, current solutions often rely on transformations that compromise the interpretability of count-based data. More broadly, there remains a need for spatially-aware, biologically-grounded integration frameworks that can mitigate sample-level variation without distorting underlying spatial structure.

As mSRT datasets continue to scale in size and complexity, computational efficiency will become increasingly critical. Popari’s spatial downsampling module offers one strategy to reduce input dimensionality while preserving biological signal by leveraging redundancy among neighboring spatial entities. Future work could explore adaptive downsampling or selective information retention, building on ideas from data-efficient machine learning to identify minimal yet sufficient representations of spatial structure [53]. In parallel, the increasing adoption of spatial multiomic technologies calls for models that can capture cross-modal interactions. While Popari can support multimodal input through simple concatenation, future extensions should more explicitly model relationships between modalities – such as transcriptional, proteomic, and epigenomic layers – to preserve interpretability and mechanistic relevance.

The growing availability of mSRT datasets across systems, conditions, and diseases will continue to raise demand for general-purpose yet interpretable tools for comparative spatial analysis. Popari addresses this gap by jointly learning metagenes through a non-negative factor model and modeling variation in their spatial affinities across samples. This structured yet flexible formulation supports both discovery and hypothesis-driven analysis, enabling deeper insights into the spatial organization of tissues in health and disease. Overall, Popari provides a unified, interpretable, and modular framework for modeling sample-to-sample spatial variation in SRT data. Its ability to capture spatial dynamics across resolution scales and biological contexts – while remaining computationally tractable and biologically meaningful – positions it as a valuable foundation for the next generation of comparative spatial omics analysis.

## Methods

### The Popari model formulation

#### Joint matrix factorization for multisample SRT

An SRT sample is represented by a tuple *D* = {*Y, C*}, where 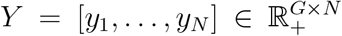 contains the gene expression profiles of *N* spatial entities (cells or spots), and 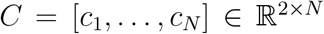 represents their 2D Euclidean coordinates, where *G* is the number of genes captured and the matrices *Y* and *C* are written in terms of column vectors. Following the non-negative matrix factorization (NMF) framework [54], *Y* can be decomposed into a factor matrix 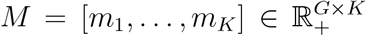, with each *m*_*k*_ ∈ 𝕊_*G−*1_, and a weight matrix 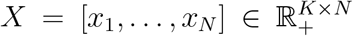. Here, *K* is the number of underlying factors (i.e., metagenes) and 𝕊_*G−*1_ is the (*G* − 1)-dimensional simplex. Constraining each column *m*_*k*_ of *M* to the simplex resolves the scaling ambiguity between *M* and *X*, enhancing interpretability. An additive noise term *E* = [*e*, …, *e*] ℝ^*G×N*^ captures unexplained variation, with each column 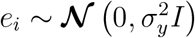, drawn i.i.d. from a multivariate Gaussian with shared variance parameter 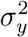.

In the multisample setting, we extend this model to accommodate *T* distinct SRT samples, where each sample *D*^*t*^ = {*Y* ^*t*^, *C*^*t*^} consists of *N* ^*t*^ entities with gene expression profiles 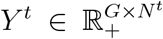 and spatial coordinates 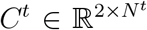. Popari assumes that the gene features are shared across all samples. For each sample, the matrix factorization model can be written as:

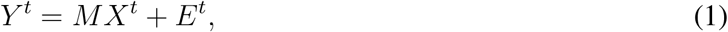

where 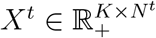 and 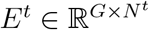 ; each column 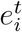 of *E*^*t*^ is drawn from a sample-specific distribution, i.e. 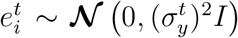. Notably, although the parameters *X*^*t*^ and 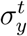 are unique to each sample, the metagenes *M* are shared across all replicates. We refer to this as the *joint NMF* formulation.

#### Probabilistic graphical model formulation for multiple samples

Popari generalizes the single-sample probabilistic graphical model introduced in the NMF-HMRF framework [17]. For each sample *t* represented by *D*^*t*^, the spatial structure is encoded as a graph **𝒢**^*t*^ = {**𝒱**^*t*^, **ℰ**^*t*^}, where vertices **𝒱**^*t*^ represent spatial entities and edges **ℰ**^*t*^ capture spatial proximity. The edge set is constructed via Delaunay triangulation of the coordinates *C*^*t*^, followed by distance thresholding (see **Supplemental Information** for details). Each vertex *i* is associated with a gene expression column vector 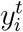 and a latent embedding vector 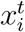. These vertex-specific quantities contribute to the likelihood function via the node potential function *ϕ*, while for each edge (*i, j*), the latent embedding vectors 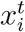 and 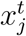 influence the likelihood through the edge potential *φ*.

The node potential is defined as:

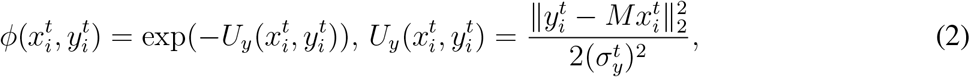

which is inversely related to the NMF reconstruction error. The edge potential is defined as:

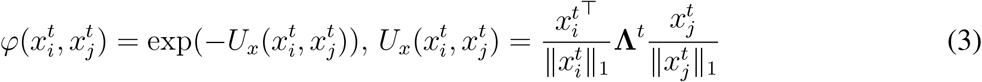

which measures the spatial accordance between embeddings of neighboring entities. Here, **Λ**^*t*^ ∈ ℝ^*K×K*^ is a matrix of learnable parameters that reflects the pairwise spatial affinities of the *K* learned metagenes. Intuitively, **Λ**^*t*^ reweights embedding dimensions of neighboring entities before computing their dot product.

By the Hammersley-Clifford theorem, the conditional likelihood of the graphical model for each sample can be expressed as the product of node and edge potentials:

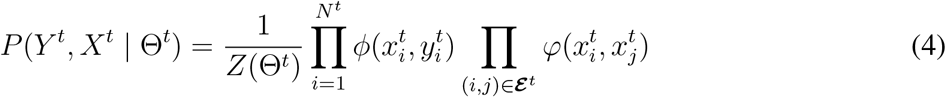

where 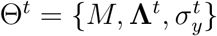 denotes the set of learnable parameters, and *Z* is the partition function ensuring proper normalization of the probability distribution.

#### Modeling multisample variation in spatial structure with probabilistic priors

Crucially, for each sample *t*, Popari estimates a distinct *K* × *K* spatial affinity matrix **Λ**^*t*^ between metagenes. This allows the model to compare and contrast the spatial structure of the transcriptome between samples.

To effectively learn **Λ**^*t*^, we introduce two regularization functions as priors. First, we place a soft constraint on the magnitude of **Λ**^*t*^ using a Gaussian prior

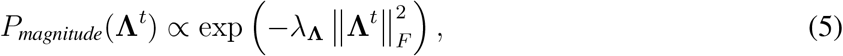

where *λ*_**Λ**_ is a hyperparameter that controls the influence of spatial information on model inference.

To incorporate sample-specific graphical models, we also introduce a differential prior on **Λ**^*t*^. We define the *average spatial affinity* 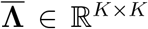, where 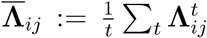, and regularize the difference 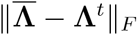. This soft constraint can also be expressed as a Gaussian prior:

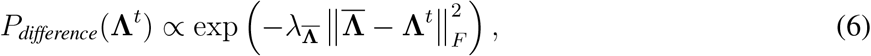

where 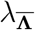 is a hyperparameter that controls the strength of differential regularization.

This prior regularizes the learned sample-specific variation in spatial structure, preventing unpenalized differences in 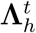 that could obscure comparisons between samples. Balancing these priors via hyperparameters *λ*_**Λ**_ and 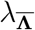 is crucial to the successful application of Popari in various settings.

Overall, the prior on the parameters is given by *P* (**Λ**^*t*^) = *P*_*magnitude*_(**Λ**^*t*^) · *P*_*difference*_(**Λ**^*t*^).

#### Multiscale hierarchical design

For each sample, Popari represents SRT data at multiple resolution levels. The original data constitutes the base resolution level, and subsequent levels are constructed by “binning” the data from the previous resolution.

Consider an SRT sample *D*_0_ = {*Y*_0_, *C*_0_}, where 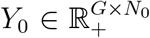 and 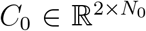. Given a set of low-resolution bins at coordinates 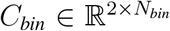, the binary *bin assignment* matrix 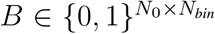 is the one-hot encoding mapping from the original coordinates *C*_0_ to the nearest coordinates *C*_bin_. The *binned expression* is given by:

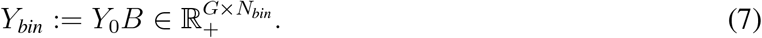

Note that *B* is a one-to-one mapping from the original entities to the low-resolution bins. This down-sampling reduces the number of spatial entities from *N*_0_ to *N*_*bin*_.

We recursively define sequential *hierarchical levels*: for each level *h*, with bin assignment matrices 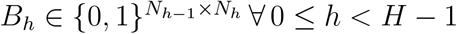, the binned expression is:

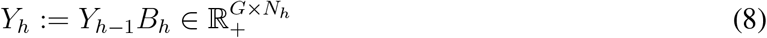

where *H* is the number of downsampling levels used. Bin coordinates 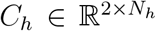 are computed as the average of the coordinates of the corresponding entities at the previous level; see **Supplemental Information** for different strategies for generating *C*_*h*_.

#### Joint hierarchical NMF-HMRF in Popari

Building on the above, we extend the NMF-HMRF to derive an overall likelihood function across all samples.

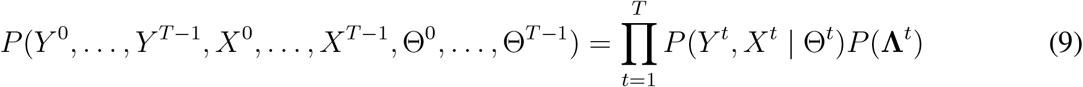

We refer to this as the *joint NMF-HMRF* model.

Each quantity involved in the joint NMF-HMRF likelihood (i.e. *M, X*^*t*^, *E*^*t*^, *Y* ^*t*^, **𝒱**^*t*^, **ℰ**^*t*^, 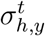, **Λ**^*t*^, and 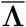) can be made specific to a hierarchical level by appending *h* to the subscript index. We notate the conditional likelihood of each sample by 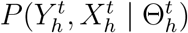, analogous to the non-hierarchical case.

Here, the weight matrix 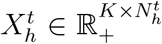, additive noise term 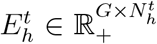, and affinity matrix 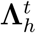 are specific to each sample *t* and level *h* pair (*t, h*), while the factor matrix *M*_*h*_ is for each level *h*.

#### Efficient optimization with alternating coordinate descent

Directly maximizing the likelihood of the joint hierarchical NMF-HMRF is computationally intractable. Instead, we use an effective approach to estimate each graph’s parameters 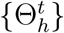 and latent states 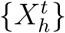. Specifically, we apply alternating coordinate descent to iteratively solve for embeddings and parameters.

The latent embeddings 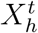 are optimized via the MAP estimate of a reparameterization of their posterior distribution, resulting in a constrained optimization problem solvable by projected coordinate descent with acceleration.

We divide the parameters into two groups: (1) differential parameters (i.e., 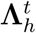 and 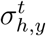) specific to each graph, and (2) shared parameters (i.e., *M*) common across all samples. The differential parameters are estimated separately for each graph. The MAP estimate of 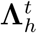 is convex and solved using the Adam optimizer, while 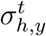 has a closed-form solution. The MAP estimate of the shared metagenes *M*_*h*_ is a strongly convex, constrained quadratic program, solved via projected gradient descent with Nesterov acceleration. See **Supplemental Information** for detailed derivations of the Popari optimization procedures.

### Details of the analysis of the multisample spatial transcriptomics datasets

For details of the preprocessing and hyperparameter selection for the application of Popari to the below datasets, see **Supplemental Information**.

#### Analysis of the STARmap PLUS dataset

For visualization of known cell types in the Popari embedding space, we first z-score normalized every latent dimension of the embeddings 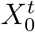 across the cell dimension (separately for each dataset). We then merged all the samples and applied the UMAP embedding with the `min_dist` parameter set to 0.05, using default settings for all other parameters. Cells were colored by their annotated labels from the original study [32].

To associate metagenes with cell types, we calculated the average value of each embedding dimension within each cell type. We then standardized the results by subtracting the minimum value per metagene and dividing by the maximum value, yielding values between 0 and 1 for each metagene. For downstream analyses, we merged the datasets that were replicates of the same condition (i.e., same timepoint and disease state). To merge the 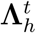 from multiple replicates, we took their elementwise average. The individual 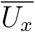 score values (i.e., the “accordance scores”; see **Supplemental Information** for the precise definition of 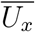) for oligodendrocytes and the 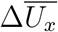 matrix were computed at level *h* = 0. The Δ**Λ**_*h*_ scores and the correlation of A*B* plaques and *p-tau* tangles with metagene expressions were computed at level *h* = 1.

To convert the raw protein images into bin-level expression values, we first divided each pixel by the maximum pixel value. We then applied a sigmoid-contrast adjustment function to each pixel value *x*:

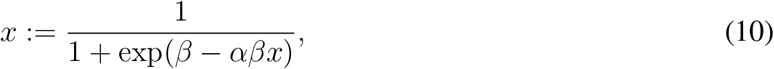

where *α* = 10 and *β* = 0.3. Finally, we scaled and discretized the pixel values to the range [0, 2^8^ − 1], and applied SquidPy’s `calculate_image_features` function to compute the protein expression score for the bins at level *h* = 1.

For spatial domain identification, we follow the postprocessing pipeline described in the **Supplemental Information**, using *D* = 13 and omitting batch effect correction. To select spatial domain marker genes for each cluster, we calculated the mean expression of each log-transformed gene value for each domain and computed the z-score across all domains. We then selected the top *n* = 10 genes with the highest z-score for each domain, and designated them as domain marker genes.

#### Analysis of the Slide-TCR-seq dataset

For spatial domain identification, we began with the embeddings at level *h* = 1 and we followed the pipeline outlined in **Supplemental Information**, with a target of *D* = 10 spatial domains and using the optional Scanorama integration step. We then applied hierarchical clustering to generate a dendrogram of the clusters. Based on this dendrogram, we manually merged the most similar clusters until only four remained, and we labeled these spatial domains according to their relative localization within the sample.

For the broad grouped-cell type analysis, we grouped the fine-grained cell types from the single-cell reference into several broader cell type categories (**Table** S1). To associate metagenes with these categories (both broad and fine-grained), we calculated the average value of each embedding dimension in each category for the level *h* = 0 embeddings. These values were thenwstandardized by subtracting the minimum and dividing by the maximum per metagene, yielding values between 0 and 1.

To assess the relationship between TRBV usage and metagene expression, we first aggregated the usage counts of all TRBV regions into a single value for each spatial entity at level *h* = 1. We then performed a permutation test comparing metagene expression in spots using TRBV versus control spots. Specifically, for each timepoint, we randomly sampled the same number of spots as those with TRBV usage and calculated their average metagene expression. This process was repeated *n* = 100 times to plot the 5th and 95th largest values to provide a 90% confidence interval on the value of the average expression for each timepoint. Metagenes exceeding the upper bound across all timepoints were deemed significantly enriched in spots with TRBV region usage.

#### Analysis of the CosMx dataset

For visualization of known cell types in the Popari embedding space, we first z-score normalized each latent dimension of the embeddings 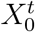 across the cell dimension (separately for each dataset).

To compute the metagene-gene set overlap AUROC, we used marker gene sets and differentially expressed gene sets defined in the original study, corresponding to broad cell types and malignant-associated cell subtypes, respectively [51]; see **Supplemental Information** for details.

To perform hierarchical clustering of samples based on spatial structure, we computed the correlation distance:

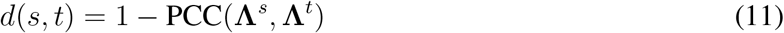

where PCC denotes the Pearson correlation coefficient, and the distance is computed between the flattened 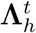 for every pair of samples *s* and *t*,

For the differential metagene proportion analysis, we calculated the *proportional metagene difference* for each single cell. For a pair of metagenes *a* and *b*, the proportional difference is given by:

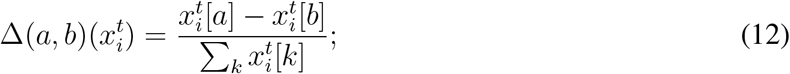

We calculated this statistic for malignant cells using metagenes m1 and m8, and for fibroblasts using metagenes m0 and m15.

## Acknowledgment

This work was supported, in part, by National Institutes of Health Common Fund 4D Nucleome Program grant UM1HG011593 (J.M.); National Institutes of Health Common Fund Cellular Senescence Network Program grant UH3CA268202 (J.M.); and National Institutes of Health grants R01HG007352 (J.M.), R01HG012303 (J.M.), R21DA061481 (J.M.), and U24HG012070 (J.M.). F.C. received funding from the Impetus Awards. J.M. was additionally supported by the Ray and Stephanie Lane Professorship, a Guggenheim Fellowship from the John Simon Guggenheim Memorial Foundation, a Google Research Award, and a Single-Cell Biology Data Insights award from the Chan Zuckerberg Initiative. S.A. was supported by the NIH Training Grant T32EB009403 (2023–2024). The funders had no role in study design, data collection and analysis, decision to publish or preparation of the manuscript.

## Code Availability

The source code for Popari can be accessed at GitHub (https://github.com/alam-shahul/popari) as well as PyPI (https://pypi.org/project/popari/).

## Author Contributions

Conceptualization, S.A., J.M.; Methodology, S.A., J.M.; Software, S.A. Investigation, S.A., T.Z., E.H., B.C., S.L., F.C., J.M.; Writing, S.A., J.M.; Funding Acquisition, F.C., J.M.

