## Supplemental Information for "Popari: Modeling multisample variation in spatial transcriptomics"

**Additional method details for the analysis of the STARmap PLUS dataset** **21**

**Additional method details for the analysis of the Slide-TCR-seq dataset** **21**

Additional method details for the analysis of the CosMx dataset 22

#### Additional details of the POPARI model

Recall that the overall likelihood of the joint hierarchical NMF-HMRF is given by:

$$P(Y, X, \Theta) = \prod_{h=1}^{H}\prod_{t=1}^{T} P(Y_h^t, X_h^t, \Theta_h^t) \tag{13}$$

$$= \prod_{h=1}^{H}\prod_{t=1}^{T} P(Y_h^t, X_h^t \mid \Theta_h^t) P(\Theta_h^t) = \prod_{h=1}^{H}\prod_{t=1}^{T} P(Y_h^t, X_h^t \mid \Theta_h^t) P(\mathbf{\Lambda}_h^t) \tag{14}$$

where $Y$, $X$, and $\Theta$ represent the joint collections of the transcriptomic data inputs, latent states, and model parameters, respectively.

It is computationally intractable to directly maximize the full likelihood function of the joint hierarchical NMF-HMRF. Instead, we simplify estimation of each graph's parameters $\{\Theta_h^t\}$ and latent states $\{X_h^t\}$ via several heuristic approximations. The first is to use the pseudo-likelihood [1], which decomposes the likelihood function over hidden states into the product of conditional likelihood for each node's hidden state, given those of its neighbors:

$$P(X_h^t \mid \Theta_h^t) \approx \prod_{i \in \boldsymbol{\mathcal{V}}_h^t} P(x_{h,i}^t \mid x_{h,\eta(i)}^t, \Theta_h^t) \tag{15}$$

where $\eta(i) = \{j \in \boldsymbol{\mathcal{V}}_h^t \mid (i,j) \in \boldsymbol{\mathcal{E}}_h^t\}$ denotes the neighbors of node $i$ in the graph, and $x_{h,\eta(i)}^t = \{x_{h,j}^t \mid j \in \eta(i)\}$ are the hidden states of the neighboring nodes.

##### Derivation of latent state optimization procedure

For each sample $t$ and hierarchical level $h$, we estimate the latent embeddings $X_h^t$ by optimizing their posterior distribution. The MAP estimate is given by:

$$\begin{aligned}\widehat{X}_h^t &= \underset{X_h^t \in \mathbb{R}_+^{K \times N_h^t}}{\operatorname{argmax}} P(X_h^t \mid Y_h^t, \Theta_h^t) = \underset{X_h^t \in \mathbb{R}_+^{K \times N_h^t}}{\operatorname{argmax}} P(Y_h^t, X_h^t \mid \Theta_h^t) \\ &= \underset{X_h^t \in \mathbb{R}_+^{K \times N_h^t}}{\operatorname{argmax}} \log P(Y_h^t, X_h^t \mid \Theta_h^t) = \underset{X_h^t \in \mathbb{R}_+^{K \times N_h^t}}{\operatorname{argmin}} \sum_{i \in \boldsymbol{\mathcal{V}}_h^t} U_y(x_{h,i}^t, y_{h,i}^t) + \sum_{(i,j) \in \boldsymbol{\mathcal{E}}_h^t} U_x(x_{h,i}^t, x_{h,j}^t)\end{aligned} \tag{16}$$

We solve this constrained optimization problem heuristically by reparameterization and projected coordinate descent with acceleration. We rewrite and expand Eqn. 16,

$$\widehat{X}^t_{h,\text{MAP}} = \underset{X^t_h \in \mathbb{R}_+^{K\times N^t_h}}{\operatorname{argmax}} \left\{ P(X^t_h \mid Y, \Theta) \right\} = \underset{X^t_h \in \mathbb{R}_+^{K\times N^t_h}}{\operatorname{argmax}} \log P(Y^t_h, X^t_h | \Theta^t_h) \tag{17}$$

$$= \underset{X^t_h \in \mathbb{R}_+^{K\times N^t_h}}{\operatorname{argmin}} \sum_{i\in\boldsymbol{\mathcal{V}}^t_h} \log \phi(x^t_{h,i}, y^t_{h,i}) + \sum_{(i,j)\in\boldsymbol{\mathcal{E}}^t_h} \log \varphi(x^t_{h,i}, x^t_{h,j}) \tag{18}$$

$$= \underset{X^t_h \in \mathbb{R}_+^{K\times N^t_h}}{\operatorname{argmin}} \left\{ \sum_{i\in\boldsymbol{\mathcal{V}}^t_h} \left[ \frac{\|y^t_{h,i} - M x^t_{h,i}\|_2^2}{2(\sigma^t_{h,y})^2} \right] + \sum_{(i,j)\in\boldsymbol{\mathcal{E}}^t_h} \frac{{x^t_{h,i}}^\top}{\|x^t_{h,i}\|_1} \boldsymbol{\Lambda}^t_h \frac{x^t_{h,j}}{\|x^t_{h,j}\|_1} \right\}. \tag{19}$$

In the last equation, we substitute $\phi$ and $\varphi$ with their definition in Eqn. 2, and 3.

For each $i$, we decompose $x^t_{h,i}$ into $s_i z_i$, where $s_i \in \mathbb{R}_+$ is a size factor representing the $L_1$ norm of $x$ and $z^t_{h,i} \in \mathbb{S}_{K-1}$ is a point on the $K$-dimensional simplex:

$$x^t_{h,i} = s^t_{h,i} z^t_{h,i} \quad \text{s.t.} \quad s^t_{h,i} \in \mathbb{R}_+, z^t_{h,i} \in \mathbb{S}_{K-1}. \tag{20}$$

Then the optimization problem becomes:

$$\underset{\forall i,\ s_i \in \mathbb{R}_+, z_i \in \mathbb{S}_{K-1}}{\operatorname{argmin}} \left\{ \left[ \frac{\|y^t_{h,i} - M z^t_{h,i} s^t_{h,i}\|_2^2}{2(\sigma^t_{h,y})^2} \right] + \sum_{(i,j)\in\boldsymbol{\mathcal{E}}^t_h} {z^t_{h,i}}^\top \boldsymbol{\Lambda}^t_h z^t_{h,j} \right\} \tag{21}$$

for a particular node $i$. We note that this problem is not necessarily convex, as the Hessian with respect to $\{s^t_{h,i}, z^t_{h,i}\}$ is not always positive semi-definite. Thus, we solve this constrained optimization heuristically using a variation of coordinate descent [2, 3], wherein we alternately update the values of either $s_i$ or $z_i$ while fixing the other set of variables.

For the update to $s^t_{h,i}$ with fixed $z^t_{h,i} = \widehat{z}^t_{h,i}$, the problem reduces to a convex, constrained, and univariate optimization problem. Thus, the closed form solution can be found by solving for the point where the derivative is 0, followed by projection to $\mathbb{R}_+$:

$$\frac{\partial}{\partial s^t_{h,i}} \left( \left[ \frac{\|y^t_{h,i} - M \widehat{z}^t_{h,i} s^t_{h,i}\|_2^2}{2(\sigma^t_{h,y})^2} \right] \right) = \frac{({\widehat{z}^t_{h,i}}^\top (M^\top M) \widehat{z}^t_{h,i}) s^t_{h,i} - {y^t_{h,i}}^\top M \widehat{z}^t_{h,i}}{(\sigma^t_{h,y})^2} \tag{22}$$

$$\widehat{s}^t_{h,i} = \max \left\{ \frac{{y^t_{h,i}}^\top M \widehat{z}^t_{h,i}}{{\widehat{z}^t_{h,i}}^\top (M^\top M) \widehat{z}^t_{h,i}}, 0 \right\} \tag{23}$$

For the update to $z^t_{h,i}$ with fixed $s^t_{h,i} = \widehat{s}^t_{h,i}$, the problem is $\alpha$-strongly convex, with the value of $\alpha$ given by the maximum eigenvalue of the Hessian:

$$\frac{\partial^2}{\partial^2 z^t_{h,i}} \left( \left[ \frac{\|y^t_{h,i} - M z^t_{h,i} s^t_{h,i}\|_2^2}{2(\sigma^t_{h,y})^2} \right] + \sum_{(i,j)\in\boldsymbol{\mathcal{E}}^t_h} {z^t_{h,i}}^\top \boldsymbol{\Lambda}^t_h z^t_{h,j} \right) = \left( \frac{s^t_{h,i}}{\sigma^t_{h,y}} \right)^2 (M^\top M) \succ 0 \tag{24}$$

Due to the requirement that $z_{h,i}^t$ lies in the $K$-dimensional simplex, we use a proximal descent method for optimization. We leverage the strong convexity of this subproblem and apply proximal Nesterov's Accelerated Gradient (NAG) method to solve it. For the proximal step after each iteration of NAG, we use Lagrange multipliers and Newton's method to obtain an approximate projection onto the simplex.

##### Derivation of optimization of model parameters

The MAP estimate of the parameters is given by:

$$\widehat{\Theta} = \underset{\Theta}{\operatorname{argmax}}\, P(Y, X, \Theta) = \underset{\Theta}{\operatorname{argmax}} \prod_{h=1}^{H} \prod_{t=1}^{T} P(Y_h^t, X_h^t \mid \Theta_h^t) P(\boldsymbol{\Lambda}_h^t) \tag{25}$$

$$= \underset{\Theta_h^t}{\operatorname{argmax}} \sum_{h=1}^{H} \sum_{t=1}^{T} \log P(Y_h^t, X_h^t \mid \Theta_h^t) + \log P(\boldsymbol{\Lambda}_h^t) \tag{26}$$

$$= \underset{\Theta_h^t}{\operatorname{argmax}} \sum_{h=1}^{H} \sum_{t=1}^{T} \sum_{i \in \boldsymbol{\mathcal{V}}_h^t} \left\{ \left[ -U_y(y_{h,i}^t, x_{h,i}^t) \right] - \sum_{(i,j) \in \boldsymbol{\mathcal{E}}_h^t} U_x(x_{h,i}^t, x_{h,j}^t) - \log Z(\Theta_h^t) + \log P(\boldsymbol{\Lambda}_h^t) \right\} \tag{27}$$

$$\approx \underset{\Theta_h^t}{\operatorname{argmax}} \sum_{h=1}^{H} \sum_{t=1}^{T} \sum_{i \in \boldsymbol{\mathcal{V}}_h^t} \left\{ \left[ -U_y(y_{h,i}^t, x_{h,i}^t) - \log Z_i(\Theta_h^t) \right] - \sum_{(i,j) \in \boldsymbol{\mathcal{E}}_h^t} U_x(x_{h,i}^t, x_{h,j}^t) + \log P(\boldsymbol{\Lambda}_h^t) \right\} \tag{28}$$

where $Z(\Theta_h^t)$ is the normalizing partition function for the conditional likelihood $P(Y_h^t, X_h^t \mid \Theta_h^t)$. The approximation in Eqn. 28 uses the mean-field assumption [4], which asserts that the global partition function $Z(\Theta_h^t)$ can be approximated as a product of per-node partition functions $Z_i(\Theta_h^t)$; that is, it assumes:

$$P(Y_h^t, X_h^t \mid \Theta_h^t) \approx \prod_{i \in \boldsymbol{\mathcal{V}}_h^t} P(x_{h,i}^t, y_{h,i}^t \mid \Theta_h^t, X_{-i}, Y_{-i}) = \prod_{i \in \boldsymbol{\mathcal{V}}_h^t} P(x_{h,i}^t, y_{h,i}^t \mid \Theta_h^t, X_{-i}) \tag{29}$$

where

$$P(x_{h,i}^t, y_{h,i}^t \mid \Theta_h^t, X_{-i}) = \frac{1}{Z_i(\Theta_h^t)} \phi\left(x_{h,i}^t, y_{h,i}^t\right) \prod_{j \in \eta(i)} \varphi\left(x_{h,i}^t, x_{h,j}^t\right) \tag{30}$$

For a particular sample $t$ and hierarchical level $h$, the MAP estimate is given by:

$$\widehat{\Theta}_h^t = \underset{\widehat{\Theta}_h^t}{\operatorname{argmax}}\, P(Y_h^t, X_h^t \mid \Theta_h^t) P(\boldsymbol{\Lambda}_h^t) = \underset{\widehat{\Theta}_h^t}{\operatorname{argmax}} \left\{ \log P(Y_h^t, X_h^t \mid \Theta_h^t) + \log P(\boldsymbol{\Lambda}_h^t) \right\} \tag{31}$$

$$\approx \underset{\Theta_h^t}{\operatorname{argmax}} \left\{ \sum_{i \in \boldsymbol{\mathcal{V}}_h^t} \left[ -U_y(y_{h,i}^t, x_{h,i}^t) - \log Z_i(\Theta_h^t) \right] - \sum_{(i,j) \in \boldsymbol{\mathcal{E}}_h^t} U_x(x_{h,i}^t, x_{h,j}^t) + \log P(\boldsymbol{\Lambda}_h^t) \right\} \tag{32}$$

$$= \underset{\Theta_h^t}{\operatorname{argmax}} \left\{ -\lambda_\Lambda \|\Lambda_h^t\|_F^2 - \lambda_{\overline{\Lambda}} \|\Lambda_h^t - \overline{\Lambda}_h\|_F^2 + \sum_{i \in \boldsymbol{\mathcal{V}}_h^t} \left[ -U_y(y_{h,i}^t, x_{h,i}^t) - \log Z_i(\Theta_h^t) \right] - \sum_{(i,j) \in \boldsymbol{\mathcal{E}}_h^t} U_x(x_{h,i}^t, x_{h,j}^t) \right\} \tag{33}$$

##### *Derivation of analytic form for partition function*

From the mean-field assumption, the per-node partition function is given by:

$$Z_i(\Theta_h^t) = \int_{x_{h,i}^t \in \mathbb{R}_+^K, y_{h,i}^t \in \mathbb{R}^G} \phi(x_{h,i}^t, y_{h,i}^t) \prod_{j \in \eta(i)} \varphi(x_{h,i}^t, x_{h,j}^t) \, \mathrm{d}x_{h,i}^t \, \mathrm{d}y_{h,i}^t \tag{34}$$

$$= \int_{x_{h,i}^t \in \mathbb{R}_+^K} \prod_{j \in \eta(i)} \varphi(x_{h,i}^t, x_{h,j}^t) \, \mathrm{d}x_{h,i}^t \int_{y_{h,i}^t \in \mathbb{R}^G} \phi(x_{h,i}^t, y_{h,i}^t) \, \mathrm{d}y_{h,i}^t \tag{35}$$

The inner integral is a multivariate Gaussian integral:

$$Z_i^Y(\Theta_h^t) := \int_{y_{h,i}^t \in \mathbb{R}^G} \phi(x_{h,i}^t, y_{h,i}^t) \, \mathrm{d}y_{h,i}^t = \int_{y_{h,i}^t \in \mathbb{R}^G} \exp\left[ -\frac{\|y_{h,i}^t - M x_{h,i}^t\|_2^2}{2\left(\sigma_{h,i}^t\right)^2} \right] \mathrm{d}y_{h,i}^t = (2\pi(\sigma_{h,i}^t)^2)^{G/2} \tag{36}$$

which is independent of $x_{h,i}^t$.

We can then simplify the partition function as:

$$Z_i(\Theta_h^t) = Z_i^Y(\Theta_h^t) \cdot \int_{x_{h,i}^t \in \mathbb{R}_+^K} \prod_{j \in \eta(i)} \varphi(x_{h,i}^t, x_{h,j}^t) \, \mathrm{d}x_{h,i}^t \tag{37}$$

Next, to simplify the remaining portion of the partition function, we again decompose $x_{h,i}^t$ into $s_i z_i$, where $s_i \in \mathbb{R}_+$ is a size factor representing the $L_1$ norm of $x$ and $z_{h,i}^t \in \mathbb{S}_{K-1}$ is a point on the $K$-dimensional simplex:

$$x_{h,i}^t = s_{h,i}^t z_{h,i}^t \quad \text{s.t.} \quad s_{h,i}^t \in \mathbb{R}_+, z_{h,i}^t \in \mathbb{S}_{K-1}. \tag{38}$$

The Jacobian of this transformation is given by:

$$\left|\frac{\partial x^t_{h,i}}{\partial(s_{h,i}, z_{h,i})}\right| = \begin{vmatrix} \frac{\partial x^t_{h,i1}}{\partial s^t_{h,i}} & \frac{\partial x^t_{h,i1}}{\partial z^t_{h,i1}} & \cdots & \frac{\partial x^t_{h,i1}}{z^t_{h,i(K-1)}} \\ \vdots & \vdots & \ddots & \vdots \\ \frac{\partial x^t_{h,i(K-1)}}{\partial s^t_{h,i}} & \frac{\partial x^t_{h,i(K-1)}}{\partial z^t_{h,i1}} & \cdots & \frac{\partial x^t_{h,i(K-1)}}{\partial z^t_{h,i(K-1)}} \\ \frac{\partial x^t_{h,iK}}{\partial s^t_{h,i}} & \frac{\partial x^t_{h,iK}}{\partial z^t_{h,i1}} & \cdots & \frac{\partial x^t_{h,iK}}{\partial z^t_{h,i(K-1)}} \end{vmatrix} = \begin{vmatrix} z^t_{h,i1} & s^t_{h,i} & \cdots & 0 \\ \vdots & \vdots & \ddots & \vdots \\ z^t_{h,i(K-1)} & 0 & \cdots & s^t_{h,i} \\ 1-\sum_{k=1}^{K-1} z^t_{h,ik} & -s^t_{h,i} & \cdots & -s^t_{h,i} \end{vmatrix} = \left(s^t_{h,i}\right)^{K-1} \tag{39}$$

Thus, the remaining integral becomes:

$$Z^X_i(\Theta^t_h) := \int_{x^t_{h,i}\in\mathbb{R}^K_+} \prod_{j\in\eta(i)} \varphi(x^t_{h,i}, x^t_{h,j})\, \mathrm{d}x^t_{h,i} = \int_{x^t_{h,i}\in\mathbb{R}^K_+} \exp\left(-\sum_{j\in\eta(i)} \frac{(x^t_{h,i})^\top}{\|x^t_{h,i}\|_2^2} \mathbf{\Lambda}^t_h \frac{x^t_{h,j}}{\|x^t_{h,j}\|_2^2}\right) \mathrm{d}x^t_{h,i} \tag{40}$$

$$= \left(\int_{s^t_{h,i}\in\mathbb{R}_+} \left(s^t_{h,i}\right)^{K-1} \mathrm{d}s^t_{h,i}\right)\left(\int_{z^t_{h,i}\in\mathbb{S}_{K-1}} \exp\left(-\sum_{j\in\eta(i)} (z^t_{h,i})^\top \mathbf{\Lambda}^t_h z^t_{h,j}\right) \mathrm{d}z^t_{h,i(K-1)} \cdots \mathrm{d}z^t_{h,i1}\right) \tag{41}$$

Because POPARI parameters do not appear in the integral over $s^t_{h,i}$, we focus on deriving an analytic form for the integral over $z^t_{h,i}$. We define $\eta^t_{h,i} = \mathbf{\Lambda}^t_h \sum_{j\in\eta(i)} z^t_{h,j}$. Furthermore, for each dimension of $\eta^t_{h,i}$, we define the function $f^t_{h,ik} : z^t_{h,ik} \mapsto \exp\left(-\eta^t_{h,ik} \cdot z^t_{h,ik}\right)$. Then, we can rewrite:

$$Z^Z_i(\Theta^t_h) := \int_{z^t_{h,i}\in\mathbb{S}_{K-1}} \exp\left(-\sum_{j\in\eta(i)} (z^t_{h,i})^\top \mathbf{\Lambda}^t_h z^t_{h,j}\right) \mathrm{d}z^t_{h,i(K-1)} \cdots \mathrm{d}z^t_{h,i1} \tag{42}$$

$$= \int_{z^t_{h,i}\in\mathbb{S}_{K-1}} \exp\left(-\left(z^t_{h,i}\right)^\top \eta^t_{h,i}\right) \mathrm{d}z^t_{h,i(K-1)} \cdots \mathrm{d}z^t_{h,i1} \tag{43}$$

$$= \int_0^q \cdots \int_0^{q-\sum_{k=1}^{K-2} z^t_{h,ik}} \exp\left(-\left(z^t_{h,i}\right)^\top \eta^t_{h,i}\right) \mathrm{d}z^t_{h,i(K-1)} \cdots \mathrm{d}z^t_{h,i1}\Big|_{q=1} \tag{44}$$

$$= \left(*_{k=1}^K f^t_{h,ik}\right)(1) \tag{45}$$

where the $*$ symbol denotes the convolution operator; that is, the integral can be expressed as the convolution of the $f$ functions evaluated at $1$.

Applying the convolution theorem in conjunction with the Laplace transform $\boldsymbol{\mathcal{L}}$:

$$\boldsymbol{\mathcal{L}}\left[\left(*_{k=1}^K f^t_{h,ik}\right)(1)\right] = \prod_{k=1}^K \boldsymbol{\mathcal{L}}[f^t_{h,ik}] = s \mapsto \prod_{k=1}^K \frac{1}{s+\eta^t_{ik}} \tag{46}$$

Using partial fraction decomposition:

$$\prod_{k=1}^{K} \frac{1}{s+\eta_{ik}^{t}} = \sum_{k=1}^{K} \frac{a_k \prod_{j\neq k}(s+\eta_{ij}^{t})}{s+\eta_{ik}^{t}} \tag{47}$$

$$\implies a_k \cdot \prod_{j\neq k}(-\eta_{ik}^{t}+\eta_{ij}^{t}) = 1 \tag{48}$$

$$\implies \prod_{k=1}^{K} \frac{1}{s+\eta_{ik}^{t}} = \sum_{k=1}^{K} \frac{1}{(s+\eta_{ik}^{t})\prod_{j\neq k}(\eta_{ij}^{t}-\eta_{ik}^{t})} \tag{49}$$

Now, since the inverse Laplace transform $\boldsymbol{\mathcal{L}}^{-1}$ is linear, we can easily compute:

$$\boldsymbol{\mathcal{L}}^{-1}\left[\boldsymbol{\mathcal{L}}\left[\left(*_{k=1}^{K} f_{h,ik}^{t}\right)(1)\right]\right] = \sum_{k=1}^{K} \frac{1}{\prod_{j\neq k}(\eta_{ij}^{t}-\eta_{ik}^{t})} \cdot \boldsymbol{\mathcal{L}}^{-1}\left[\frac{1}{(s+\eta_{ik}^{t})}\right] \tag{50}$$

$$= \sum_{k=1}^{K} \frac{\exp\left(-\eta_{ik}^{t}\right)}{\prod_{j\neq k}(\eta_{ij}^{t}-\eta_{ik}^{t})} \tag{51}$$

This final analytic form can be computed efficiently in each iteration of the parameter update steps. Because it requires the multiplication of many small values, we use a signed version of the "log-sum-exp" trick to compute it in a numerically stable manner.

##### *Optimization of* $M$

The parameter $M$ appears only in the $U_y(y_{h,i}^t, x_{h,i}^t)$ terms. Thus, it is obtained by solving the following strongly convex constrained optimization problem:

$$\underset{M\in\mathbb{S}_{G-1}^{K}}{\operatorname{argmax}} \sum_{i\in\boldsymbol{\mathcal{V}}_h^t} -U_y(y_{h,i}^t, x_{h,i}^t) = \underset{M\in\mathbb{S}_{G-1}^{K}}{\operatorname{argmin}} \sum_{i\in\boldsymbol{\mathcal{V}}_h^t} \|y_{h,i}^t - Mx_{h,i}^t\|_2^2 \tag{52}$$

We solve this in each iteration using projected NAG descent, with the maximum possible step size $\frac{1}{\lambda}$ that guarantees convergence, where $\lambda$ is the maximum eigenvalue of the Hessian of the objective.

##### *Optimization of* $\sigma_{h,y}^t$

The partial derivatives of terms related to $(\sigma_{h,y}^t)^{-1}$ are:

$$\frac{\partial}{\partial(\sigma_{h,y}^t)^{-1}} \log\phi(x_{h,i}^t, y_{h,i}^t) = -(\sigma_{h,y}^t)^{-1}\|y_{h,i}^t - Mx_{h,i}^t\|_2^2 \quad \frac{\partial}{\partial(\sigma_{h,y}^t)^{-1}} \log Z_i^Y(\Theta_h^t) = -\frac{G}{(\sigma_{h,y}^t)^{-1}} \tag{53}$$

This implies that the function is overall concave and can be maximized by solving for the point at which the partial derivatives are zero. This yields the closed-form update:

$$\sigma_{h,y}^t = \sqrt{\frac{1}{N_h^t G} \sum_{i\in\boldsymbol{\mathcal{V}}_h^t} \|y_{h,i}^t - Mx_{h,i}^t\|_2^2} \tag{54}$$

##### *Optimization of* $\mathbf{\Lambda}_h^t$

In the objective function, $\mathbf{\Lambda}_h^t$ appears in the regularization term $\log P(\mathbf{\Lambda_h^t})$, the linear term $U_x(x_{h,i}^t, x_{h,j}^t)$ and the partition function component $Z_i^Z(\Theta_h^t)$. Thus, we minimize the following:

$$\lambda_\Lambda \|\Lambda_h^t\|_F^2 + \lambda_{\overline{\Lambda}} \|\Lambda_h^t - \overline{\Lambda}_h\|_F^2 + \sum_{i \in \mathcal{V}_h^t} \sum_{j \in \eta i} U_x(x_{h,i}^t, x_{h,j}^t) + \sum_{i \in \mathcal{V}_h^t} Z_i^Z(\Theta_h^t) \tag{55}$$

using the Adam optimizer [5].

#### Adjacency matrix construction

We use Squidpy's [6] `spatial_neighbors` function to determine the adjacency matrix for each graph $\mathcal{G}_h^t$. We first call the function with `delaunay=True` to compute the canonical Delaunay triangulation. However, from experience, we find that the the distribution of edge lengths from the canonical Delaunay triangulation tends to be right-skewed, with a number of long edges unlikely to reflect true spatial interactions in tissue. To address this, we filter out excessively long edges by computing a `cutoff` value at the $94.5^{\text{th}}$ percentile of all edge lengths. We then call `spatial_neighbors` again with `delaunay=True` and `radius=[0, cutoff]`, ensuring that abnormally long edges are excluded from the final adjacency matrix. The advantage of this approach is that it is fully automated, as the `cutoff` parameter is determined statistically rather than relying on prior biological knowledge,

#### Generation of bin assignments

The hierarchical formulation of POPARI is valid for any bin assignment $B_h \in \{0,1\}^{N_{h-1} \times N_h}$ that satisfies two constraints: (1) each column sums to 1, and (2) $N_{h-1} < N_h$. Accordingly, there are multiple reasonable strategies for generating the bin assignment matrix. We present two approaches here, each parametrized by a desired downsampling rate $r_{\text{bin}}$:

1. *Grid-based downsampling*. A 2D grid of equally-spaced bins is generated to span the ranges of the $x$ and $y$ coordinates for the input coordinates. The bin spacing is chosen heuristically to ensure $N_{h-1}$ is close to $r_{\text{bin}} \cdot N_h$. Each spatial entity from the original resolution is then assigned to its nearest bin based on Euclidean distance.

2. *Partition-based downsampling*. Using the spatial graph constructed during the POPARI graphical model setup, the graph is partitioned into $N_{h-1} = \lfloor r_{\text{bin}} \cdot N_h \rfloor$ components of approximately equal size. This is accomplished using `pymetis`, a wrapper for the METIS graph-partitioning software [7].

#### Initialization of model parameters at lowest resolution

The results of the POPARI optimization procedure are sensitive to the initial values of the latent states and model parameters, making good initialization crucial for producing interpretable output. To initialize the metagene matrix and latent states, we first concatenate the preprocessed gene expression matrices after binning at the lowest resolution level:

$$Y_{cat} := [Y^0_{H-1}, Y^1_{H-1}, \ldots, Y^{T-1}_{H-1}] \tag{56}$$

Next, we use principal component analysis (PCA) to obtain a reduced-dimensional representation of each entity's gene expression. We then apply Leiden clustering on the PCA embeddings to obtain exactly $K$ clusters, where $K$ is the number of metagenes in the metagene matrix $M$. To obtain such a clustering, we binary search for the Leiden clustering resolution parameter that yields exactly $K$ clusters in the original expression space. We initialize the columns of $M$ as the centroids of the $K$ clusters, and the embeddings of the spatial entities $X_{cat}$ are the one-hot encodings of the cluster assignments. Splitting $X_{cat}$ gives the initial embeddings for each sample at the lowest resolution:

$$[X^0_{H-1}, X^1_{H-1}, \ldots, X^{T-1}_{H-1}] := X_{cat} \tag{57}$$

Because the higher-resolution embeddings are not used during the core optimization steps, these are initialized randomly, which greatly reduces the initialization wall time.

The spatial affinity matrices $\mathbf{\Lambda}^t_h$ are initialized using the results of the Leiden clustering. For each graph $\boldsymbol{\mathcal{G}}^t_h = \{\boldsymbol{\mathcal{V}}^t_h, \boldsymbol{\mathcal{E}}^t_h\}$, we calculate the empirical spatial correlation matrix as described below in the "Additional general method details" section.

Following this static initialization, model training is further warm-started. Specifically, the $X^t_H$, $M$ and $\sigma^t_{H,y}$ parameters are optimized for a small number of "non-spatial preiterations" using the standard non-negative matrix factorization (NMF) objective [8]; during this stage, $\mathbf{\Lambda}^t_h$ remains fixed.

Optionally, in a second warm-start phase, the $X^t_H$, $M$, $\sigma^t_{H,y}$ and $\sigma^t_{H,y}$ are further optimized through a few "spatial preiterations" using the SPICEMIX objective [9], where the sample-specific $\mathbf{\Lambda}^t_H$ are simplified into a single spatial affinity matrix. Finally, the POPARI optimization procedure is carried out using these initialized values.

##### Hierarchical propagation of $\mathbf{\Lambda}^t$ parameters and initialization of $X^t_h$ at higher resolution levels

To efficiently optimize the POPARI parameters across different resolutions, we use the learned $\mathbf{\Lambda}^t$ parameters and $X^t_h$ embeddings from lower resolutions to initialize the optimization for higher resolutions.

Specifically, we introduce a new optimization formulation that leverages the hierarchical structure of the data. Given the embeddings $X^t_h = \widehat{X}^t_h$ and the bin assignments $B^t_h$ for a sample $t$ and hierarchical level $h$, we define $X^t_{h-1}$ as the solution to the following problem:

$$\widehat{X}^t_{h-1} = \underset{X^t_{h-1} \in \mathbb{R}_+^{K \times N^t_h}}{\operatorname{argmin}} \frac{1}{2} \left( \|Y^t_{h-1} - MX^t_{h-1}\|^2_F + \|\widehat{X}^t_h - X^t_{h-1}B^t_h\|^2_F \right) \tag{58}$$

Intuitively, this formulation replaces the spatial component of the embedding objective function with a soft constraint enforcing that, for each low-resolution bin, the embeddings of its assigned high-resolution entities sum up to its own embedding, i.e. $X^t_h = X^t_{h-1}B^t_h$. Because the low-resolution bin embeddings were learned using the original spatially-informed latent embedding objective, this soft constraint is sufficient to ensure that the high-resolution embeddings inherit spatial information. The above objective is a constrained convex optimization problem, which we solve efficiently using projected gradient descent.

In particular, the derivative of the objective with respect to $X_{h-1}^t$ is given by:

$$\frac{\partial}{\partial X_{h-1}^t}\left(\frac{1}{2}\|Y_{h-1}^t - MX_{h-1}^t\|_F^2 + \frac{1}{2}\|\widehat{X}_h^t - X_{h-1}^t B_h^t\|_F^2\right) \tag{59}$$

$$= M^\top M X_{h-1}^t - (Y_{h-1}^t)^\top M + X_{h-1}^t (B_h^t)^\top B_h^t (B_h^t)^\top - B_h^t \left(\widehat{X}_{h-1}^t\right)^\top \tag{60}$$

For all datasets, rather than using PyTorch's `autograd` module, we explicitly compute this gradient and optimize using alternating iterations of Adam with projection to $\mathbb{R}_+^{K\times N_h^t}$ until convergence, thereby obtaining the superresolved embeddings.

After initializing $X_h^t$ at higher resolutions using this procedure, we complete parameter and embedding optimization using just a few iterations of the POPARI optimization. In practice, we find that 10 iterations are sufficient.

##### Grid search for hyperparameter selection with MLflow

To enable organized sweeping of POPARI hyperparameter combinations, we use MLflow [10], an open-source framework for tracking and visualizing the results of machine learning experiments. MLflow enables parallelization of hyperparameter grid searches across multiple GPUs.

Leveraging MLflow, we provide a script to grid search for the best choice of POPARI hyperparameters, including the number of metagenes $K$, the number of hierarchical levels $H$, the desired downsampling rate at each hierarchical level $r_{\text{bin}}$, the spatial affinity magnitude penalty $\lambda_{\mathbf{\Lambda}}$ and the spatial affinity difference penalty $\lambda_{\overline{\mathbf{\Lambda}}}$. After completion of the runs, the MLflow GUI is used to qualitatively compare runs and select the best hyperparameter combination for downstream analysis.

#### Additional details of the simulation generative model

In flexibly model a variety of SRT contexts and generate the two simulations presented in the main text, we developed a novel multisample spatial transcriptomics simulation framework that affords precise control over generation of both gene expression and spatial organization of simulated entities. A factor-based generative model with user-specified parameters is used to generate the multisample gene expression data, while the spatial structure of the simulated SRT samples is defined using both specified parameters and a graphical user interface (GUI).

Simulation parameters controlling gene expression include: $K$, the number of ground truth metagenes; $C$, the number of ground truth cell types; $G$, the number of genes; `cell_type_definitions`, a matrix in $\mathbb{R}_+^{C\times K}$ indicating the relative importance of each metagene to each cell type; `metagene_variation_probabilities`, which controls the overlap in the gene-based definition of the ground truth metagenes; `sigX_scale` ($\sigma_x$), which controls the intra-cell type variance of the ground truth embeddings; `sigY_scale` ($\sigma_y$), which sets the variance of Gaussian noise independently added to gene expression profiles.

Parameters controlling spatial structure include: $L$, the number of layers/spatial domains in the simulation, and `layer_distributions`, a mapping from each layer to a vector in $\mathbb{S}_{C-1}$ indicating the cell type frequencies in each layer. The `grid_size` parameter specifies the number of spots in each row and column of the sample, leading to $N = \texttt{grid\_size}^2$ total spatial entities per simulated sample. The GUI is used to precisely mark ground truth spatial domains. It enables the user to mark domain "landmarks", and simulated entities are assigned to domains based on the nearest landmarks. Finally, we model structured sparsity in the output expression data using the `spatial_sparsity` parameter, which specifies a fraction of spatial entities to downsample. For each gene $g$, `spatial_sparsity` fraction of the entities are sampled from the spatial graph such that the set of sampled cells forms an independent set (i.e., there are no edges between entities in the set); the expression of gene $g$ in the sampled entities is set to $0$.

We used this framework to generate both the hierarchical and differential simulations for benchmarking against baseline methods. However, the framework is general and supports the generation of synthetic mSRT datasets with realistic yet complex structure. In particular, the GUI component greatly enhances the flexibility of the simulator for modeling arbitrary spatial patterns, enabling it to mimic both imaging-based and array-based SRT modalities.

##### Generative model for $M$

The metagenes $M$ are synthesized sequentially, one column (i.e. metagene) at a time. For the first metagene $m_0 \in \mathbb{R}^G$, the weight of each gene in the metagene is sampled from a Gamma distribution with shape parameter given by `real_metagene_parameter`. For subsequent metagenes, gene weights are similarly sampled from the same Gamma distribution, but a proportion of the gene weights are shared with the previous metagene; the proportion is specified by the vector `metagene_variation_probabilities` $\in [0, 1)^K$, where the $i$-th entry indicates the proportion of gene weights that $m_i$ should share with $m_{i-1}$.

Finally, the metagenes are then divided by their L1 norm such that they lie on the simplex, i.e. $m_k := \frac{m_k}{\|m_k\|_1}$

Note that in the multisample, differential simulation, the metagenes $M$ are shared across all samples, so the generation of $M$ is performed only once per simulation.

##### Generative model for $X^t$

For each sample $t$, $\texttt{grid\_size}^2$ spatial entities are equidistantly spaced within a $1 \times 1$ unit square. Then, the user assigns each entity to a "spatial domain." This is accomplished using a GUI input system, by which the user defines a set of landmark coordinates for each domain; entities are assigned the domain corresponding to that of the nearest landmark coordinate. The entities in each domain are randomly assigned a cell type according to the `cell_type_distributions` parameter, which contains a vector of cell type proportion for each spatial domain. Next, for each entity $i$ and for each embedding dimension $k$, the simulated embedding $x_i^t[k]$ is sampled according to a cell-type specific truncated Gaussian distribution. The mean of the distribution is given by the `cell_type_definitions` parameter, which maps from cell types to mean vectors $\mu \in \mathbb{R}_+^K$. The scale of the distribution is given by the parameter $\sigma_x$; its support is lower bounded by $0$, and upper bounded by $100 \cdot \sigma_x + \mu_k$. After generation, the embeddings $X^t$ are scaled such that each cell's embedding lies on the simplex:

$$x_i^t := \frac{x_i^t}{\|x_i^t\|_1} \tag{61}$$

To produce the final ground truth cell embeddings $X^t$, each cell $i$ is multiplied by a size factor $s_i^t$ drawn from a Gamma distribution with shape parameter $\frac{K}{\lambda_s}$ and scale parameter $\lambda_S$, i.e.

$$s_i^t \sim \text{Gamma}\left(\frac{K}{\lambda_s}, \lambda_S\right) \tag{62}$$

$$x_i^t := s_i^t \cdot x_i^t \tag{63}$$

##### Generative model for $Y^t$

To generate the output expression values $Y^t$, we first compute the intermediate $\widetilde{Y^t}$:

$$\widetilde{Y^t} := MX^t \tag{64}$$

Next, each entity's expression $y_i^t$ is sampled according to a multivariate Gaussian distribution, with mean vector given by $\widetilde{y_i^t}$ and covariance matrix given by $\sigma_y \mathbf{I}$, where $\sigma_y$ is an input parameter. Finally, the absolute value is taken element-wise to ensure that the generated expression values are nonnegative.

##### Specific simulation parameters for hierarchical simulation

For the hierarchical simulation setting, we designed a scenario with $C = 8$ cell types, $G = 100$ genes and $K = 9$ underlying metagenes. We used `grid_size` $= 36$, implying $N = 1296$ total entities. We used $L = 4$ spatial domains, which constituted vertical layers subdividing the unit square. The layers, named `L1`, `L2`, `L3` and `L4`, were $0.35$, $0.15$, $0.25$ and $0.25$ units in width, respectively. Four cell types, named $A_{\texttt{L1}}$, $A_{\texttt{L2}}$, $A_{\texttt{L3}}$ and $A_{\texttt{L4}}$, have a weighting of $0.5$ for metagene 0 and a weighting of $1$ for metagenes 5, 6, 7 and 8, respectively. Two cell types, named $B_1$ and $B_2$, have a weighting of $0.5$ for metagene 1 and a weighting of $1$ for metagenes 3 and 4, respectively. Finally, the $C_{\texttt{ubiquitous}}$ cell type has a weighting of 1 for metagene 2, whereas the $C_{\texttt{L1}}$ cell type has a weighting of 1 for metagene 2 and a weighting of $0.5$ for metagene 5. In addition, metagenes 0 and 1 are linked such that they share $70\%$ of their gene weights; similarly, metagenes 3 and 4 also share $70\%$ of their gene weights, and metagenes 5, 6, 7 and 8 share $70\%$ of their gene weights. The `sigY_scale` is 1, and the `sigX_scale` is 0.1. The `spatial_sparsity` is 0.15.

##### Specific simulation parameters for differential simulation

For the differential simulation setting, we designed a scenario with $C = 9$ cell types, $G = 100$ genes and $K = 11$ underlying metagenes. We used a `grid_size` $= 15$ per FOV, yielding $N = 225$ entities per replicate. Ten FOVs were simulated, resulting in a total of $2250$ entities across all samples. These ten

FOVs were split into two categories, $t_0$ and $t_1$, each with five replicates.

As in the hierarchical simulation, there are $L = 4$ layers named `L1`, `L2`, `L3` and `L4`, and these are $0.35$, $0.15$, $0.25$ and $0.25$ units in width, respectively.

Four cell types, named $A_{\texttt{L1}}$, $A_{\texttt{L2}}$, $A_{\texttt{L3}}$ and $A_{\texttt{L4}}$, have a weighting of $0.5$ for metagene 0 and a weighting of $1$ for metagenes 7, 8, 9 and 10, respectively.

Another four cell types, named $B_{\texttt{L1}}$, $B_{\texttt{L2}}$, $B_{\texttt{L3}}$ and $B_{\texttt{L4}}$, have a weighting of $0.5$ for metagene 1 and a weighting of $1$ for metagenes 3, 4, 5 and 6, respectively.

Finally, the $C_{\texttt{ubiquitous}}$ cell type has a weighting of $1$ for metagene 2.

The `sigY_scale` is 1, and the `sigX_scale` is 0.1.

In addition, metagenes 0 and 1 are linked such that they share $90\%$ of their gene weights. Metagenes 3, 4, 5 and 6 also share $90\%$ of their gene weights, and do metagenes 7, 8, 9 and 10.

### Additional general method details

#### Details of the $U_{x_{\text{pair}}}$ computation

To further interpret individual entity-entity spatial accordances learned by POPARI, we further analyzed the values of the $U_x$ potential function across edges of the spatial graph. Given a pair of metagenes $k, l$ and the embeddings $x_w, x_j$ for two neighboring entities $i, j$, we compute the metagene pair-specific component of the spatial accordance $U_{x_{\text{pair}}}$ as:

$$z_i = \frac{x_i}{\|x_i\|_2}$$

$$z_{i,\text{pair}} = (z_{is})_{s \in (k,l)} \tag{65}$$

$$\mathbf{\Lambda}_{\text{pair}} = (\mathbf{\Lambda}_{st})_{s,t \in (k,l)} \tag{66}$$

$$U_{x_{\text{pair}}}(x_i, x_j) = {z_{i,\text{pair}}}^\top \mathbf{\Lambda}_{\text{pair}} z_{j,\text{pair}} \tag{67}$$

That is, the $U_{x_{\text{pair}}}$ only captures the spatial interaction between the cells directly corresponding to the cross-affinity and self-affinities between the the elements of the dataset. These values are directly visualized *in situ* without further processing.

#### Details of the $\overline{U_x}$ computation

Given a set of categorical labels (e.g. cell types, spatial domains, etc.) over the spatial entities with $L$ total categories, we compute the "$\overline{U_x}$ score" vector $\iota^t_{h,i} \in \mathbb{R}^L$ for each entity $i$, which captures its average accordance with entities of each category. Recall the definition of the edge potential function:

$$U_x(x^t_{h,i}, x^t_{h,j}) = \frac{{x^t_{h,i}}^\top}{\|x^t_{h,i}\|_2} \mathbf{\Lambda}^t_h \frac{x^t_{h,j}}{\|x^t_{h,j}\|_2} \tag{68}$$

The $k$-th element of the $\iota_{h,i}^t$ is then given by:

$$\iota_{h,i}^t[k] := \begin{cases} \sum_{j\in\eta(i)} \left(\frac{\mathbb{I}[\ell(i)=k]}{\sum_{j\in\eta(i)} \mathbb{I}[\ell(j)=k]}\right) \cdot U_x(x_{h,i}^t, x_{h,j}^t) & \sum_{j\in\eta(i)} \mathbb{I}[\ell(j)=k] > 0 \\ 0 & \text{otherwise,} \end{cases} \tag{69}$$

where $\eta(i)$ is the set of neighbors of $i$ in the spatial graph $\boldsymbol{\mathcal{G}}_h^t$ and $\ell$ is the function that returns the categorical label of an entity given its index.

The $\overline{U_x}$ score vectors of all entities in graph $\boldsymbol{\mathcal{G}}_h^t$ can be summarized into an average "$\overline{U_x}$ matrix" $\in \mathbb{R}^{L\times L}$, where the entry in row $k$ and column $l$ represents the average accordance of entities in category $k$ with neighboring entities in category $l$:

$$\overline{U_x}[k,l] := \begin{cases} \sum_{i=1}^{N_h^t} \left(\frac{\mathbb{I}[\ell(i)=k]}{\sum_{i=1}^{N_h^t} \mathbb{I}[\ell(i)=k]}\right) \cdot \iota_{h,i}^t[l] & \sum_{i=1}^{N_h^t} \mathbb{I}[\ell(i)=k] > 0 \\ 0 & \text{otherwise.} \end{cases} \tag{70}$$

#### Details of the empirical spatial correlation computation

Given a spatial graph $\boldsymbol{\mathcal{G}} = \{\boldsymbol{\mathcal{V}}, \boldsymbol{\mathcal{E}}\}$ with embeddings $X \in \mathbb{R}_+^{K\times|\boldsymbol{\mathcal{V}}|}$ (where $K$ is the number of metagenes/embedding dimensions), we calculate the empirical spatial correlation matrix of embeddings over edges of the graph. In this context, we treat the edges as though they are directed, i.e. the "first" node in each edge is distinguished from the "second" node.

We define mean and standard deviation vectors for the embeddings of first and second nodes:

$$\mu_i := \sum_{(i,j)\in\boldsymbol{\mathcal{E}}} \frac{x_i}{|\boldsymbol{\mathcal{E}}|}, \;\; \sigma_i := \sqrt{\sum_{(i,j)\in\boldsymbol{\mathcal{E}}} \frac{(x_i-\mu_i)^2}{|\boldsymbol{\mathcal{E}}|}} \tag{71}$$

$$\mu_j := \sum_{(i,j)\in\boldsymbol{\mathcal{E}}} \frac{x_j}{|\boldsymbol{\mathcal{E}}|}, \;\; \sigma_j := \sqrt{\sum_{(i,j)\in\boldsymbol{\mathcal{E}}} \frac{(x_j-\mu_j)^2}{|\boldsymbol{\mathcal{E}}|}} \tag{72}$$

where each operation above is vectorized.

Using these statistics, we then compute the empirical correlation matrix $\widetilde{\boldsymbol{\Lambda}} \in \mathbb{R}^{K\times K}$ between the embeddings of the first and second nodes:

$$\widetilde{\boldsymbol{\Lambda}}_h^t := -\sum_{(i,j)\in\boldsymbol{\mathcal{E}}} \left(\frac{x_i-\mu_i}{\sigma_i}\right)\left(\frac{x_j-\mu_j}{\sigma_j}\right)^\top \tag{73}$$

Finally, we symmetrize the matrix by adding it to its transpose, and then zero-center the result by subtracting the mean value of the matrix from all entries.

#### Details of spatial domain identification pipeline

To automatically discover spatial domain clusters in each dataset, we created a postprocessing pipeline as follows. Given a target number of spatial domains $D$,

1. Z-score normalize each embedding dimension across all spatial entities, including all samples.

2. Remove outlier expression levels for each metagene by clipping all values above the 99th percentile per sample.

3. Divide the expression levels of each metagene by the sum of its expression values in the top 1% spots per sample.

4. (Optional) Use Scanorama [11] to perform batch effect correction, using the sample name as the batch.

5. Smooth metagene expression levels to the average of itself and its neighbors:

$$x_i := \frac{\sum_{j \in \eta(i) \cup \{i\}} x_j}{|\eta(i)| + 1} \tag{74}$$

   where $x_i \in \mathbb{R}^K$ is the vector-valued postprocessed embedding of spot $i$, and $\eta(i)$ represents the neighbors of spot $i$ in the graph.

6. Run Leiden clustering using the `scanpy` package's `leiden` API, with `n_neighbors=20` and `resolution=1`. If this does not yield exactly $D$ clusters, then we binary search for the Leiden clustering resolution parameter which yields exactly $D$ clusters.

7. Smooth cluster labels until convergence. Specifically, let $l_i^0$ be the raw cluster label of spot $i$ produced in the previous step, and repeat the following procedure iteratively starting with $t = 0$:

$$l_i^{t+1} := \begin{cases} \ell & \frac{1}{|\eta(i)|+1} \sum_{j \in \eta(i) \cup \{i\}} \mathbb{I}(l_j^t = \ell) > 0.5 \\ l_i^t & \text{no such label } \ell \text{ exists} \end{cases} \tag{75}$$

#### Details of metagene signature gene set selection

To summarize the most important genes in a metagene, we first rank the genes in the input in ascending order by their respective weights within the metagene.

We then use the `kneedle` algorithm [12] to pick an appropriate "elbow" rank at which to threshold the set of signature genes: all genes above this rank are considered part of the signature gene set, and all genes below this rank are excluded from the set.

We use a sensitivity parameter value of $S = 0.1$ for all applications of the `kneedle` algorithm in our analysis.

##### Details of metagene-gene set AUROC score computation

To quantify the concordance of a metagene's gene rankings to a predefined gene set of interest, we summarize the overlap of the gene set with the top-ranked genes in the metagene for every possible gene weight threshold.

That is, treating membership in the gene set as the positive label, we compute the receiver operatic characteristic (ROC) curve for all $G$ weight thresholds.

Thus, the maximum area under the ROC (AUROC) score of $1$ is achieved when all of the genes in the gene set are ranked higher within the metagene than all of the genes not in the gene set, and a score lower than $0.5$ indicates that a metagene is underenriched for the gene set (i.e., less than random chance).

#### Additional method details of the simulation evaluation

##### Details of the ground truth evaluation procedure

###### *Evaluation metrics*

We developed two metrics to evaluate performance on the simulated datasets. First, to quantify the similarity in the spatial distribution of the expression value of a ground truth metagene and an inferred metagene, we formulated the *spatial Wasserstein distance* (SWD) as the solution to a "min cost flow with demands" optimization problem.

Given a sample with $n$ simulated spatial entities, we calculate the pairwise distance matrix $C' \in \mathbb{Z}_+^{n\times n}$ based on the spatial coordinates; these constitute the costs of moving flow from one spatial entity to another.

The demands and supplies for the nodes $B' \in \mathbb{Z}^n$ is given by the difference between their ground truth embedding values and their inferred embedding values. We compute the SWD efficiently using Google's OR-Tools suite [13]; further details of the linear programming formulation are given below.

We compute the SWD between every pair of ground truth and inferred metagene embedding values, yielding the all pairs SWD matrix $D^{\text{SWD}} \in \mathbb{R}_+^{K\times K}$, where $K$ is the number of both ground truth and inferred metagenes.

To produce a single metric, we first obtain the minimum distance from each ground truth metagene to any of the inferred metagene. Then, among these minima, we select the maximum value. We denote this summary metric as the *maxmin spatial Wasserstein distance* (mSWD). Because the mSWD is a rigorously defined distance metric, it has a strict interpretation: the best (minimum) possible value is $0$, and larger values indicate the inferiority of a method in recapitulating the ground truth metagene spatial patterns.

We also formulated the *affinity Spearman correlation* (ASC) in order to capture the correlation between the learned spatial affinity matrix $\Lambda_{\text{inf}} \in \mathbb{R}^{K\times K}$ and the expected spatial affinities.

We first use the all pairs SWD matrix $D^{\text{SWD}}$ in order find the best one-to-one matching between the ground truth and inferred metagenes using Scipy's `linear_sum_assignment` function, with $D_{\text{SWD}}$

as the cost matrix. This function finds a boolean matching matrix $M \in [0,1]^{K \times K}$ that minimizes

$$\sum_{0 \le i < K} \sum_{0 \le j < K} d_{ij}^{\mathrm{SWD}} m_{ij} \tag{76}$$

where each row and column of $M$ is constrained to sum to 1.

We then compute the empirical spatial correlation $\mathbf{\Lambda}_{\text{empirical}}$ between all ground truth metagene embeddings, which gives a proxy for the expected ground truth spatial affinity; see above for details of the empirical correlation computation.

The rows and columns of $\mathbf{\Lambda}_{\text{inferred}}$ are permuted using the matching $M$, yielding $\mathbf{\Lambda}_{\text{permuted}}$, allowing for direct comparison with $\mathbf{\Lambda}_{\text{empirical}}$.

The final ASC value is given by the Spearman correlation between elements of $\mathbf{\Lambda}_{\text{permuted}}$ and $\mathbf{\Lambda}_{\text{empirical}}$. The maximum value of 1 indicates that the method is able to perfectly recapitulate the relative ranks of the expected ground truth spatial affinities.

##### *Details of baseline evaluations*

For the comparisons to NMF, hierarchical NMF (NMF-H), SpiceMix, disjoint SpiceMix (SM-D) and joint SpiceMix (SM-J), we implemented these baselines as using the Popari codebase. SpiceMix, SM-D and SM-J use exactly the same implementation and optimization scheme as Popari, except that they omit differential regularization of spatial affinities. Furthermore, for SM-D, a separate spatial affinity matrix is learned for each replicate, whereas for SM-J a single affinity matrix is learned for all replicates. For the NMF and NMF-H baselines, the optimization is similar to Popari, except that spatial information is ignored.

For the NSF [14] baseline used in the hierarchical simulation, we use the PyPI package implemented at `https://github.com/willtownes/spatial-factorization-py`. We use the hyperparameters `L=9` (the same as the choice of $K$ for Popari), `J=100` (i.e. the number of genes in the simulation), and `N=1296` (i.e. the number of spots in the simulation) for all runs. For the INSPIRE [15] baseline used in the differnetial simulation, we use the implementation available at `https://github.com/jiazhao97/INSPIRE`. We use the hyperparameters `n_spatial_factors=11` (the same as the choice of $K$ for Popari) and `n_training_steps=5000` for all runs. For all other hyperparameters, the default is used.

##### *Evaluation design*

For each simulation scenario, we generate 20 replicates of the simulation using different *random seeds* for the generative simulation framework.

In addition, because the outputs are stochastic, we run each method with 5 different *random states* for each replicate;

overall, we run each method 100 times for each simulation setting.

To summarize the evaluation for a setting, we take the best performance of each method across all random states for each replicate, i.e. for the mSWD, we take the minimum value, and for the ASC, we take the maximum value.

For the differential simulation, because there are multiple FOVs, we additionally take the average of these metrics across all FOVs. These yields the final values used in the scatterplots in **Fig.** 2B-C.

For all simulation scenarios and all methods, we set the metagene hyperparameter to the same value as the number of ground truth metagenes that are used in the generation of the simulation.

We use a value of $\lambda_{\mathbf{\Lambda}} = 10^{-4}$ for both SPICEMIX and POPARI in all simulation scenarios.

We use a value of $\lambda_{\overline{\Lambda}} = 10^{-3}$ for POPARI in the differential simulation scenario, and a binning downsample rate of $f_B = 0.2$ with 2 hierarchical levels for POPARI in the hierarchical scenario.

##### Details of the spatial Wasserstein distance (SWD) linear programming formulation

Given a sample with $n$ simulated spatial entities, we calculate the pairwise distance matrix $C \in \mathbb{R}_+^{n \times n}$ based on the spatial coordinates; self-distances along the diagonal are assigned a distance of positive infinity. The distance matrix is then scaled such that the minimum value is $10^4$, and the results are rounded down to the nearest integer:

$$c'_{ij} := \left\lfloor 10^4 \frac{c_{ij}}{\min(C)} \right\rfloor \forall\, 0 \le i, j < n \tag{77}$$

The ground truth embedding values $X \in \mathbb{R}_+^n$ and the inferred embedding values $\widehat{X} \in \mathbb{R}_+^n$ are scaled so that they sum to 1. The demands $B \in \mathbb{R}^n$ are given by the difference between the ground truth embedding value and the inferred embedding value, scaled such that the maximum demand is $10^4$, and the results are rounded down to the nearest integer:

$$b_i := x_i - \widehat{x}_i;\; b'_i := \left\lfloor 10^4 \frac{b_i}{\max(B)} \right\rfloor \forall\, 0 \le i < n \tag{78}$$

For the construction of the edge set, a directed edge is added between two spatial entities $i$ and $j$ if one has a positive demand and the other has a negative demand, that is,

$$\boldsymbol{\mathcal{E}} = \{(i,j) \forall\, 0 < i, j < n : b_i > 0, b_j < 0\} \tag{79}$$

Each edge is assigned a capacity of $10^4$, and a cost equal to the scaled spatial distance between the two nodes.

The SWD is then given as the solution to the following optimization problem:

$$\min \sum_{(i,j) \in \boldsymbol{\mathcal{E}}} c'_{ij} f_{ij} \tag{80}$$

$$\text{subject to}$$

$$\sum_{0 \le i < n} f_{ij} - \sum_{0 \le k < n} f_{jk} = b'_j \;\forall\, 0 \le j < n$$

$$0 \le f_{ij} \le 10^4 \;\forall\, (i,j) \in \boldsymbol{\mathcal{E}}$$

where $f_{ij}$ represents the "flow" that must move from entity $i$ to entity $j$ in order to balance the distributions. We use Google's OR-Tools to efficiently compute the SWD [13].

#### Additional method details for the analysis of the STARmap PLUS dataset

We used all eight available samples. We applied the `normalize_total` and `log_1p` functions, yielding the following transformation of the raw SRT gene counts $Y_{ig}$:

$$Y'_{ig} := \log\left(1 + \left(10^4 \frac{Y_{ig}}{\sum_{g'} Y_{ig'}}\right)\right) \tag{81}$$

Because the gene panel is limited, we use all 2,766 available genes for further analysis. Finally, we generate the spatial neighborhood graph using `spatial_neighbors`.

We first performed a grid search to optimize for POPARI hyperparameters, including the number of metagenes $K$, the spatial affinity regularization $\lambda_{\mathbf{\Lambda}}$, the differential regularization $\lambda_{\overline{\mathbf{\Lambda}}}$, and the downsample rate $r_{bin}$ using grid-based downsampling. Each run used 10 non-spatial preiterations, 50 spatial preiterations and 200 POPARI iterations. Based on qualitative plots, including heatmaps of the spatial affinity matrices $\mathbf{\Lambda}_h^t$ and *in situ* plots of the metagene embeddings $X_h^t$, we selected a run with $K = 20$, $\lambda_{\mathbf{\Lambda}} = 10^{-4}$, $\lambda_{\overline{\mathbf{\Lambda}}} = 10^{-4}$ and $r_{bin} = 0.2$.

#### Additional method details for the analysis of the Slide-TCR-seq dataset

From the available samples, we selected samples `day_3.1`, `week_4.2`, `week_5.2`, `week_13.1` and `week_15.1`, which correspond to mice at age 3, 28, 35, 91 and 105 days, respectively. We first manually annotated the boundary between the thymic and non-thymic region in each slide, and excluded the non-thymic spots from further analysis. We then apply Scanpy's [16] `normalize_total` and `log_1p` functions, yielding the following transformation of the raw SRT gene counts $Y_{ig}$:

$$Y'_{ig} := \log\left(1 + \left(10^4 \frac{Y_{ig}}{\sum_{g'} Y_{ig'}}\right)\right) \tag{82}$$

We used `filter_cells` on the transformed counts with `min_counts` $= 50$. To select genes, we used Scanpy's `highly_variable_genes` function with `n_top_genes` $= 500$ separately on each slide, and take the union of the HVGs for each slide; then, we use the `filter_genes` function for this set with `min_cells` $= 3$ for each slide, and take the intersection to produce the final gene set. Additionally, we call differentially expressed genes (DEGs) for each cell type from an scRNA-seq thymic cell atlas, and add these to the gene set. We also use marker genes identified in [17] and add these to the gene set. Overall, this yields a set of 6,371 genes for further analysis. Finally, we generate the spatial neighborhood graph using `spatial_neighbors`.

In a similar manner to the STARmap PLUS evaluation, we performed a grid search for the number of metagenes $K$, the spatial affinity regularization $\lambda_{\mathbf{\Lambda}}$, the differential regularization $\lambda_{\overline{\mathbf{\Lambda}}}$, and the downsample rate $r_{bin}$ using grid-based downsampling. Each run used 10 non-spatial preiterations, 50 spatial preiterations and 200 POPARI iterations. Based on qualitative plots, including heatmaps of the spatial affinity matrices $\mathbf{\Lambda}_h^t$ and *in situ* plots of the metagene embeddings $X_h^t$, we selected a run with $K = 20$, $\lambda_{\mathbf{\Lambda}} = 10^{-4}$, $\lambda_{\overline{\mathbf{\Lambda}}} = 10^{-4}$ and $r_{bin} = 0.2$.

#### Additional method details for the analysis of the CosMx dataset

We used all 27 adnexal tumor samples from the "Discovery" dataset in [18]. We applied the `normalize_total` and `log_1p` functions, yielding the following transformation of the raw SRT gene counts $Y_{ig}$:

$$Y'_{ig} := \log\left(1 + \left(10^4 \frac{Y_{ig}}{\sum_{g'} Y_{ig'}}\right)\right) \tag{83}$$

To select genes, we used Scanpy's `highly_variable_genes` function with `n_top_genes` $=$ $500$ separately on each slide, and take the union of the HVGs for each slide; then, we use the `filter_genes` function for this set with `min_cells` $= 3$ for each slide, and take the intersection to produce the final gene set. This yields 969 genes overall. Finally, we generate the spatial neighborhood graph using `spatial_neighbors`.

We first performed a grid search to optimize for POPARI hyperparameters, including the number of metagenes $K$, the spatial affinity regularization $\lambda_{\mathbf{\Lambda}}$, the differential regularization $\lambda_{\overline{\mathbf{\Lambda}}}$. Each run used 10 non-spatial preiterations and no spatial preiterations; due to runtime constraints, each run used 100 POPARI iterations. Based on qualitative plots, including heatmaps of the spatial affinity matrices $\mathbf{\Lambda}_h^t$ and *in situ* plots of the metagene embeddings $X_h^t$, we selected a run with $K = 20$, $\lambda_{\mathbf{\Lambda}} = 10^{-4}$ and $\lambda_{\overline{\mathbf{\Lambda}}} = 10^{-2}$.

### Supplementary Figures

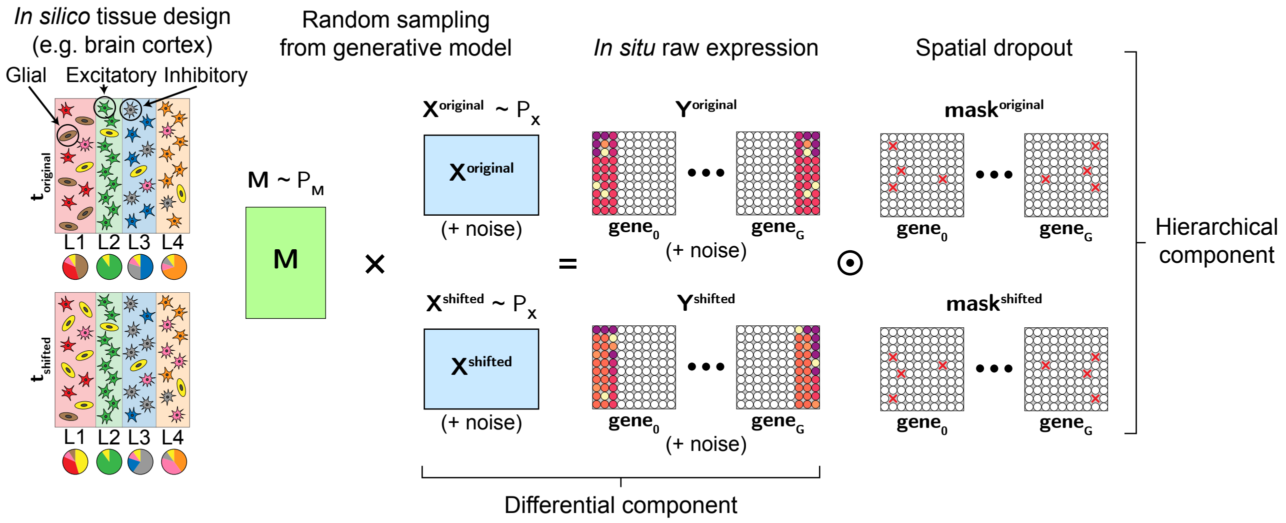

**Figure S1:** Cartoon depicting the simulation generative model. In the differential component, the pie charts indicate the relative proportions of cell types in each simulated spatial domain; these proportions differ between the $\mathbf{t}_{\text{original}}$ and $\mathbf{t}_{\text{shifted}}$ samples. In the hierarchical component, the red Xs indicate the per-gene spatial entities for which expression values are dropped out to zero. The circle with the dot in the center represents the Hadamard product of the masks with the raw simulated expression.

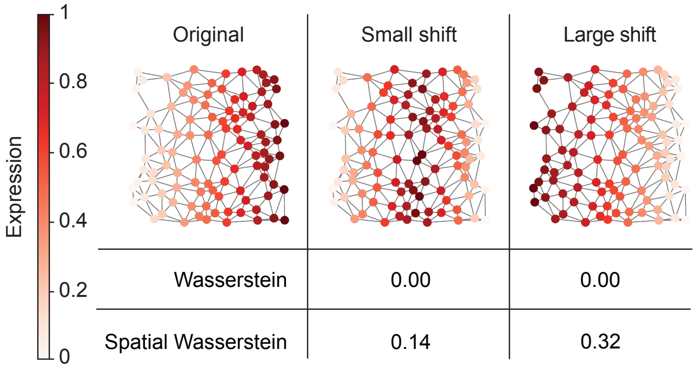

| | Original | Small shift | Large shift |
|---|---|---|---|
| Wasserstein | | 0.00 | 0.00 |
| Spatial Wasserstein | | 0.14 | 0.32 |

**Figure S2:** Comparison between the Wasserstein distance metric and the proposed Spatial Wasserstein distance (SWD) metric for toy SRT datasets. The “original” toy SRT dataset consists of 100 spatial entities, each with $xy$-coordinates randomly sampled from a $1 \times 1$ unit square and with expression value equal to the $x$-coordinate of each entity. The number in each cell of the table corresponds to the distance between the “original” dataset and the corresponding shifted dataset under the selected metric. The “large shift” toy SRT dataset was created by reversing the expression values of the “original” entities according to their $x$-coordinate. The “small shift” toy SRT dataset was created by assigning every other sorted expression value (starting with the first) to the first half of the sorted $x$-coordinates, and for the rest of the expression values reversing their order and assigning them to the second half of the sorted $x$-coordinates.

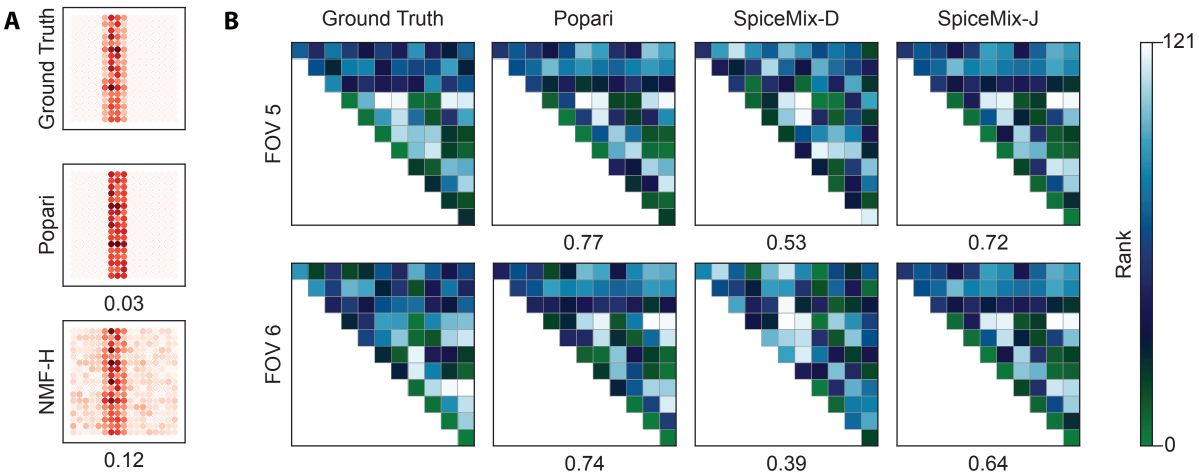

**Figure S3:** Extended visualization of representative examples for the hierarchical and differential simulations. **A**. *In situ* visualization of metagene 6 embedding values for one replicate of the hierarchical simulation at hierarchical level $h = 1$. For the ground truth, the *in situ* embedding is visualized by binning the ground truth embeddings at hierarchical level $h = 0$ using the same binning matrix employed by POPARI and NMF-H. The number below each method indicates the SWD between the binned ground truth embedding and the respective method's embedding. **B**. Ranks of the empirical/learned spatial affinities for the ground truth/benchmark methods, respectively, in FOV 5 and 6 from the differential simulation. In case of ties, elements are assigned the average rank of the tied elements. The number below each method is the absolute value of the Spearman correlation between the method's learned spatial affinity and the ground truth empirical affinity.

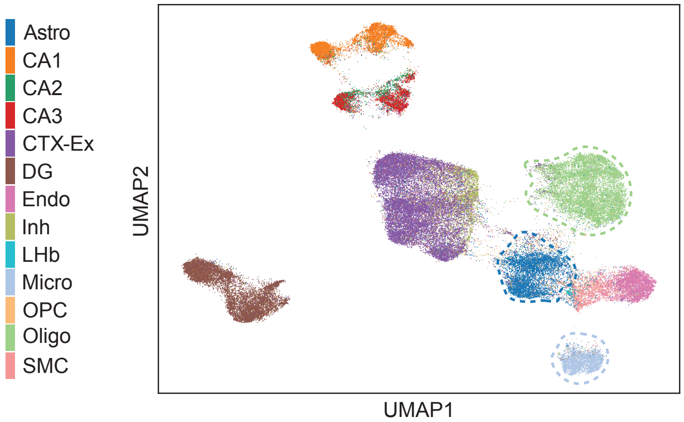

**Figure S4:** UMAP reduction of the embeddings $X^t$ at hierarchical level $h = 0$ produced by POPARI. The color key indicates the cell type identity of each cell, as labeled in the original STARmap PLUS study [19]. Cell type abbreviations: Astro: astrocytes; CA1: CA1 neurons; CA2: CA2 neurons; CA3: CA3 neurons; CTX-Ex: cortex excitatory neurons; DG: dentate gyrus neurons; Endo: endothelial cells; Inh: inhibitory neurons; LHb: LHb neurons; Micro: microglia; OPC: oligodendryocte precursor cells; Oligo: oligodendrocytes; SMC: smooth muscle cells.

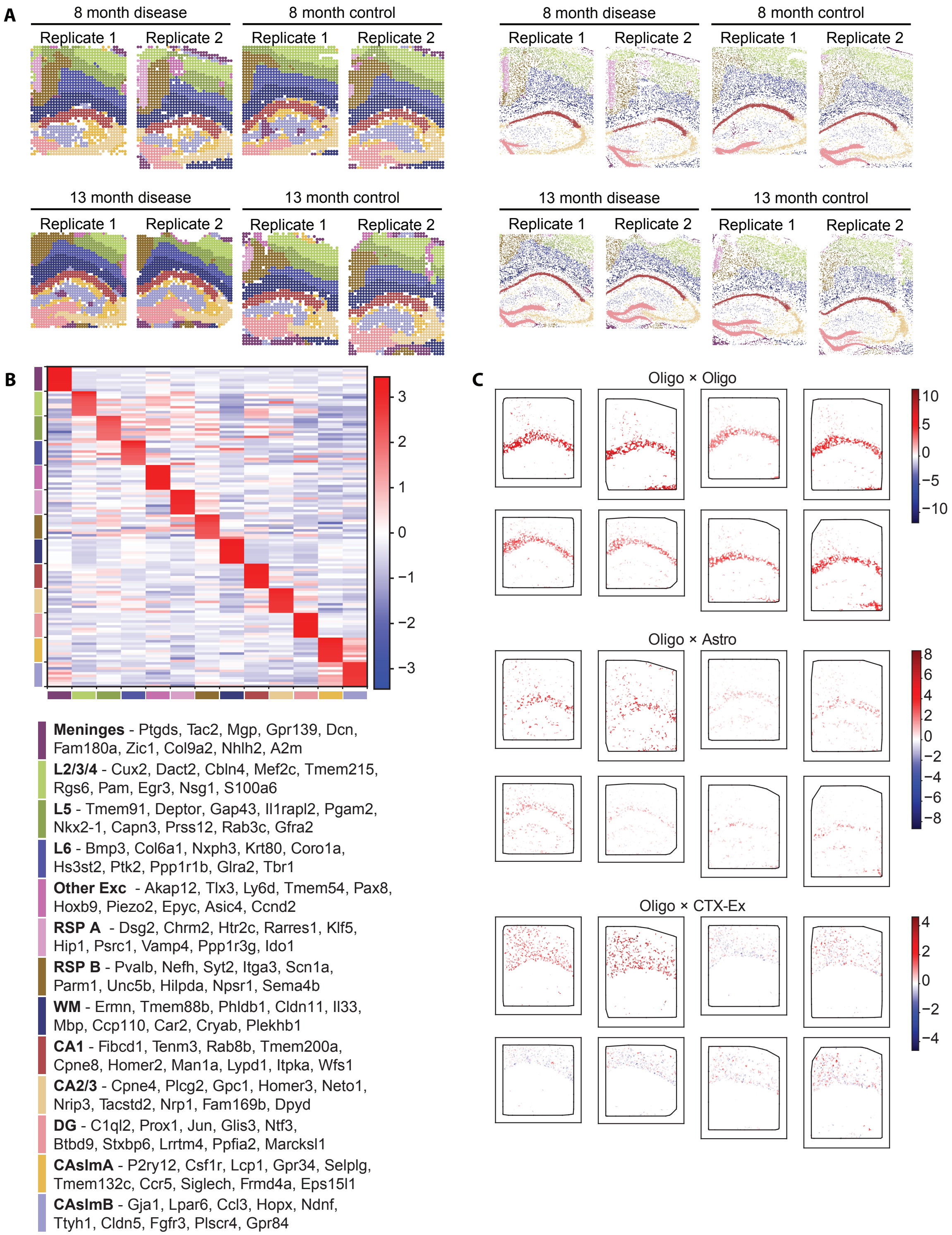

**Figure S5:** Extended interpretation of spatial domains and cell-cell interactions in STARmap PLUS dataset. **A.** (left) Spatial domain segmentation from level $h = 1$ embeddings. (right) Propagation of spatial domain labels to original spatial entities at hierarchical level $h = 0$. **B.** Average z-scored expression of marker genes across spatial domains. Every ten rows of the heatmap correspond to the top ten differential genes in a spatial domain. **C.** *In situ* plot of $\overline{U_x}$ values for oligodendrocytes with other oligodendrocytes (top), astrocytes (middle) and cortex excitatory neurons (bottom).

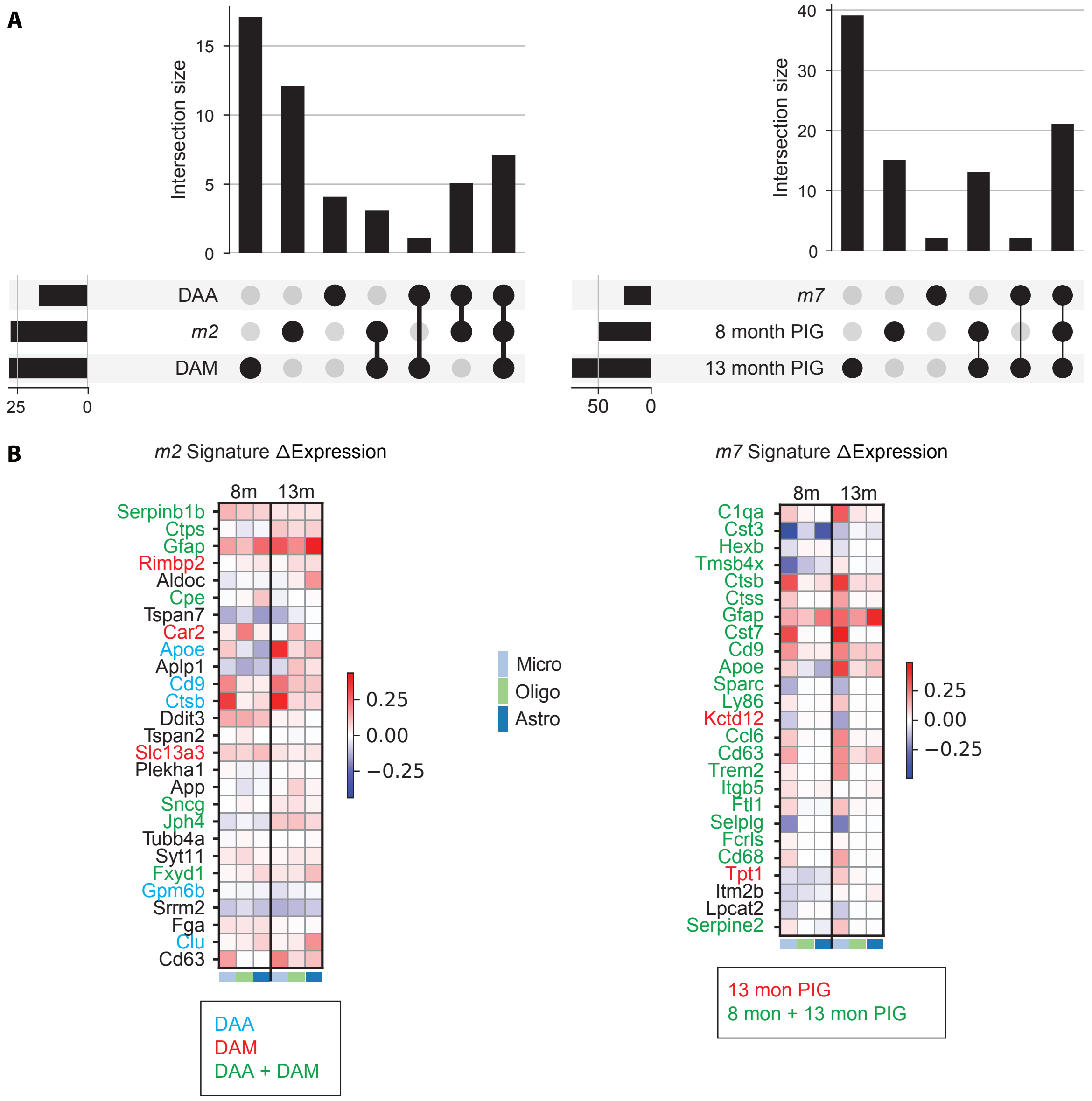

**Figure S6:** Extended interpretation of metagene identity in the STARmap PLUS dataset. **A.** Upset plots showing the overlap of disease-associated astrocyte (DAA) and disease-associated microglia (DAM) marker gene sets with the signature genes in metagene m2 (left), and the overlap of plaque-induced genes (PIGs) from 8-month and 13-month mice with metagene m7 (right). All validation gene sets were provided in the original STARmap PLUS study [19]. **B.** ΔExpression of the top signature genes across the three glial cell types – microglia, oligodendrocytes, and astrocytes – for metagenes m2 (left) and m7 (right). Colored gene names indicate membership of these signature genes in the relevant validation gene sets.

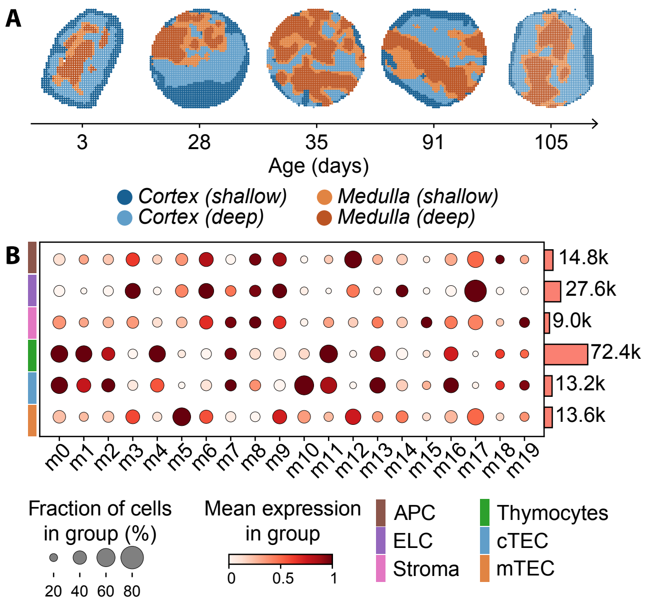

**Figure S7:** Extended interpretation of POPARI embeddings at both coarse and fine resolutions. **A.** Spatial domain identification of POPARI across the five Slide-TCR-seq thymus samples. Blue colors indicate cortex-related domains, whereas orange colors indicate medulla-related domains. **B.** Correspondence between POPARI metagenes and cell types, quantified by an enrichment plot of each metagene's mean expression value for each cell type. The marginal histogram indicates the total count of Slide-TCR-seq spots labeled with the respective cell type by RCTD.

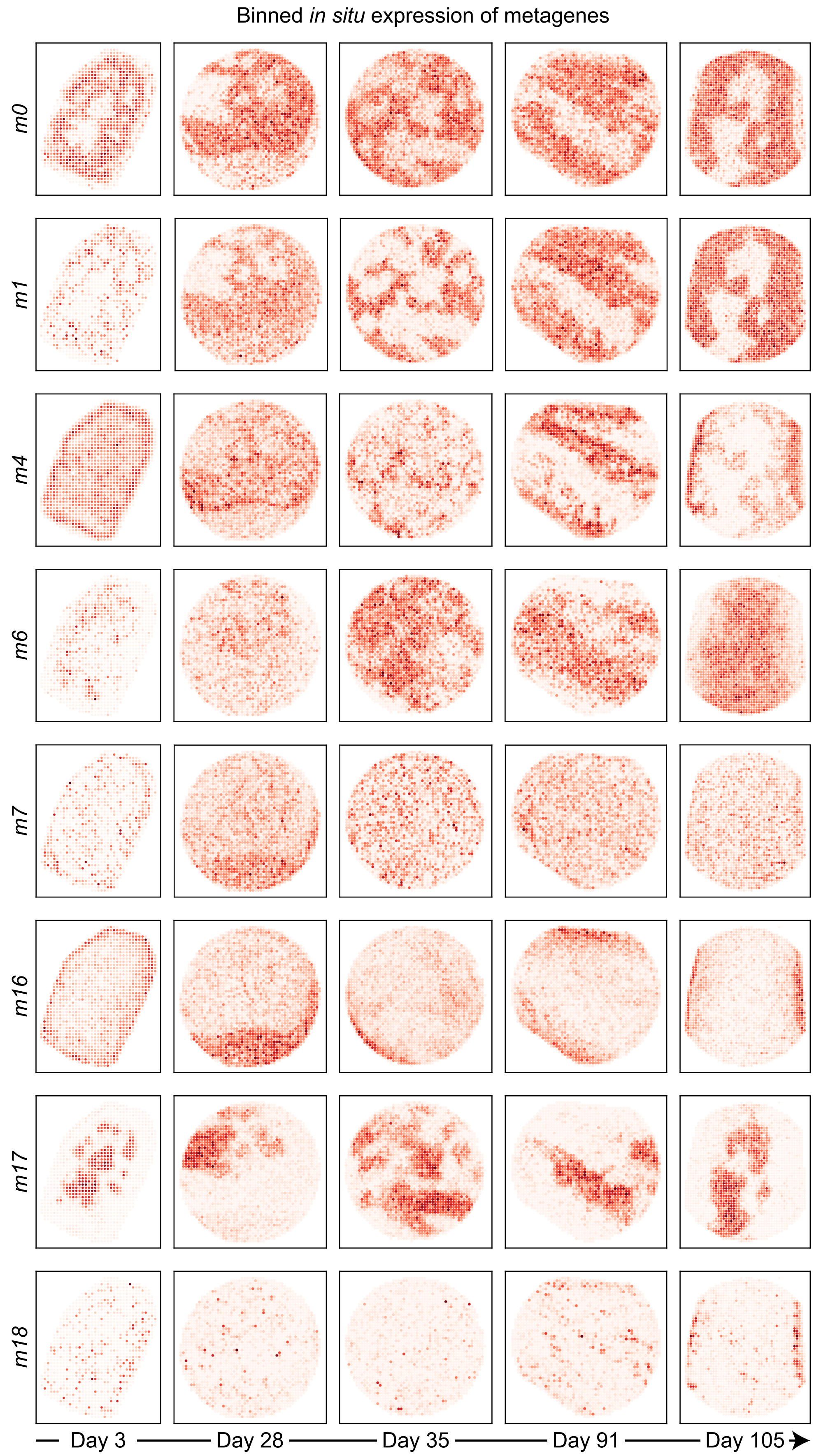

**Figure S8:** *In situ* plots of embeddings at hierarchical level $h = 1$ for all metagenes from the Slide-TCR-seq dataset.

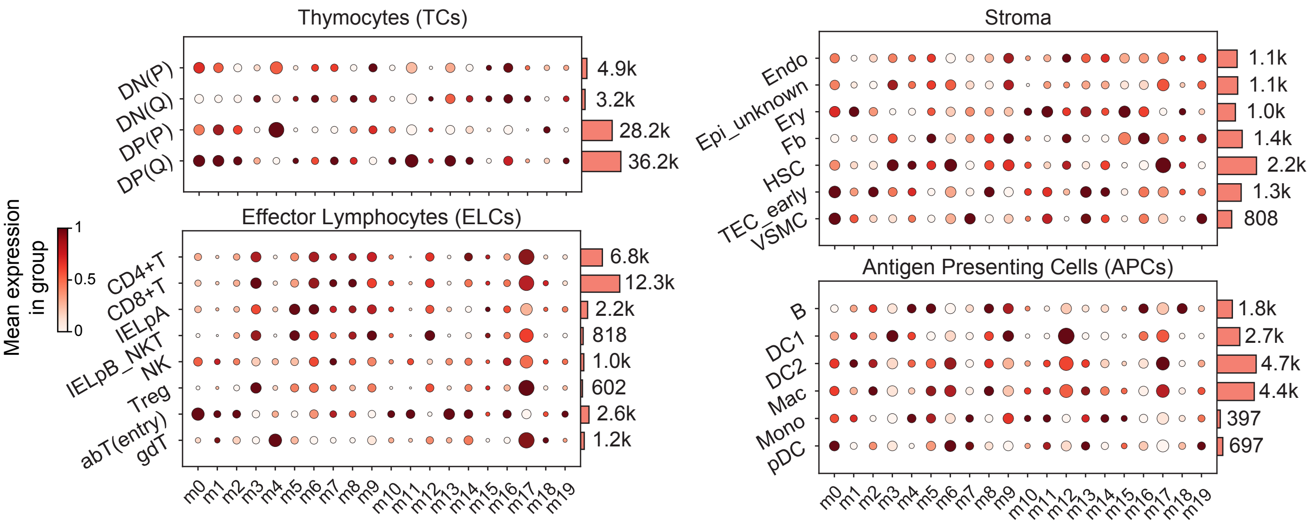

**Figure S9:** Correspondence between cell types and average metagene expression in the Slide-TCR-seq data for all fine-grained cell types from the RCTD analysis. For each of the four broad cell type categories, normalized mean expression is computed separately for the associated fine-grained cell types. The mapping from broad to fine-grained categories is given in **Table** S1.

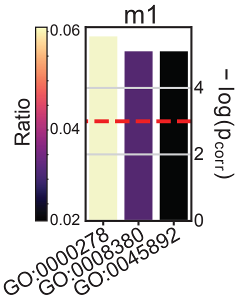

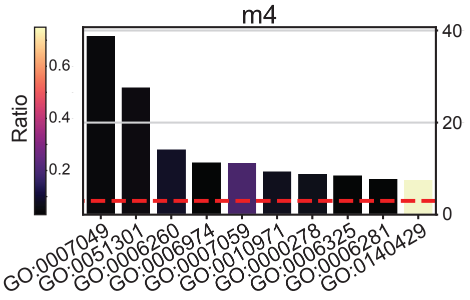

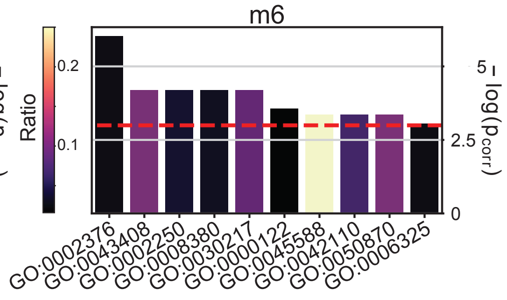

GO:0000278 - mitotic cell cycle
GO:0008380 - RNA splicing
GO:0045892 - negative regulation of DNA-templated transcription

GO:0007049 - cell cycle
GO:0051301 - cell division
GO:0006260 - DNA replication
GO:0006974 - DNA damage response
GO:0007059 - chromosome segregation
GO:0010971 - positive regulation of G2/M transition of mitotic cell cycle
GO:0000278 - mitotic cell cycle
GO:0006325 - chromatin organization
GO:0006281 - DNA repair
GO:0140429 - positive regulation of mitotic sister chromatid separation

GO:0002376 - immune system process
GO:0043408 - regulation of MAPK cascade
GO:0002250 - adaptive immune response
GO:0008380 - RNA splicing
GO:0030217 - T cell differentiation
GO:0000122 - negative regulation of transcription by RNA polymerase II
GO:0045588 - positive regulation of gamma-delta T cell differentiation
GO:0042110 - T cell activation
GO:0050870 - positive regulation of T cell activation
GO:0006325 - chromatin organization

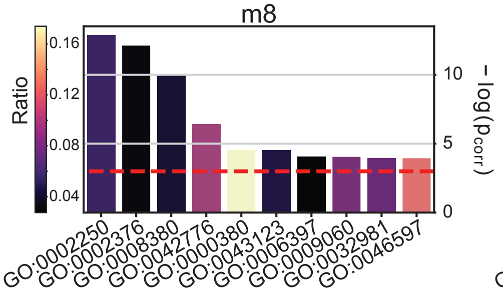

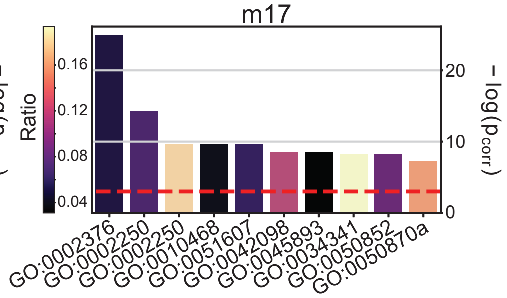

GO:0002250 - adaptive immune response
GO:0002376 - immune system process
GO:0008380 - RNA splicing
GO:0042776 - proton motive force-driven mitochondrial ATP synthesis
GO:0000380 - alternative mRNA splicing, via spliceosome
GO:0043123 - positive regulation of canonical NF-kappaB signal transduction
GO:0006397 - mRNA processing
GO:0009060 - aerobic respiration
GO:0032981 - mitochondrial respiratory chain complex I assembly
GO:0046597 - negative regulation of viral entry into host cell

GO:0002376 - immune system process
GO:0002250 - adaptive immune response
GO:0002250 - antigen processing and presentation
GO:0010468 - regulation of gene expression
GO:0051607 - defense response to virus
GO:0042098 - T cell proliferation
GO:0045893 - positive regulation of DNA-template transcription
GO:0034341 - response to type II interferon
GO:0050852 - T cell receptor signaling pathway
GO:0050870 - positive regulation of T cell activation

**Figure S10:** Gene Ontology (GO) analysis of selected spatiotemporally variable metagenes in Slide-TCR-seq dataset. The $x$-axis in each plot indicates the negative log of the FDR-corrected p-value ($p_{\text{corr}}$) for each GO term. Bar colors reflect the ratio of genes shared between the metagene signature gene set ($n_{\text{signature}}$) and the GO term gene set ($n_{\text{GO}}$). The red dashed line marks the significance threshold value of $-\log(0.05)$.

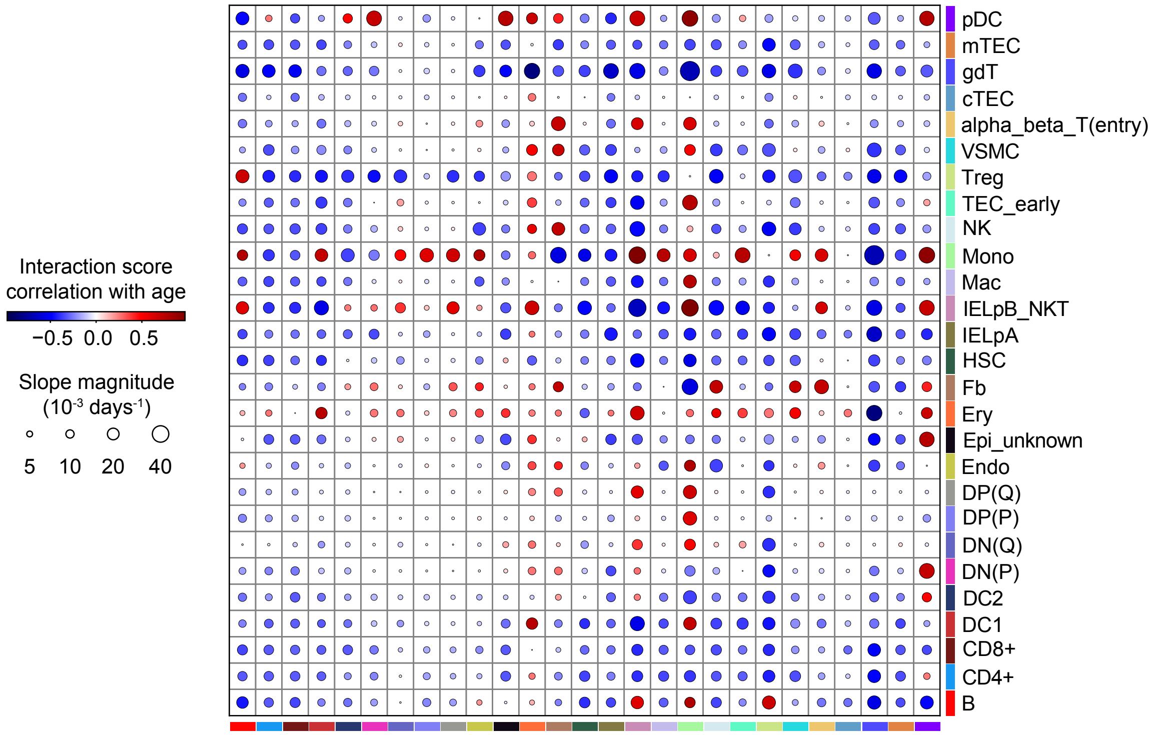

**Figure S11:** Pearson correlation and regression slope of all fine-grained cell type-cell type interaction score pairs with age in the Slide-TCR-seq data.

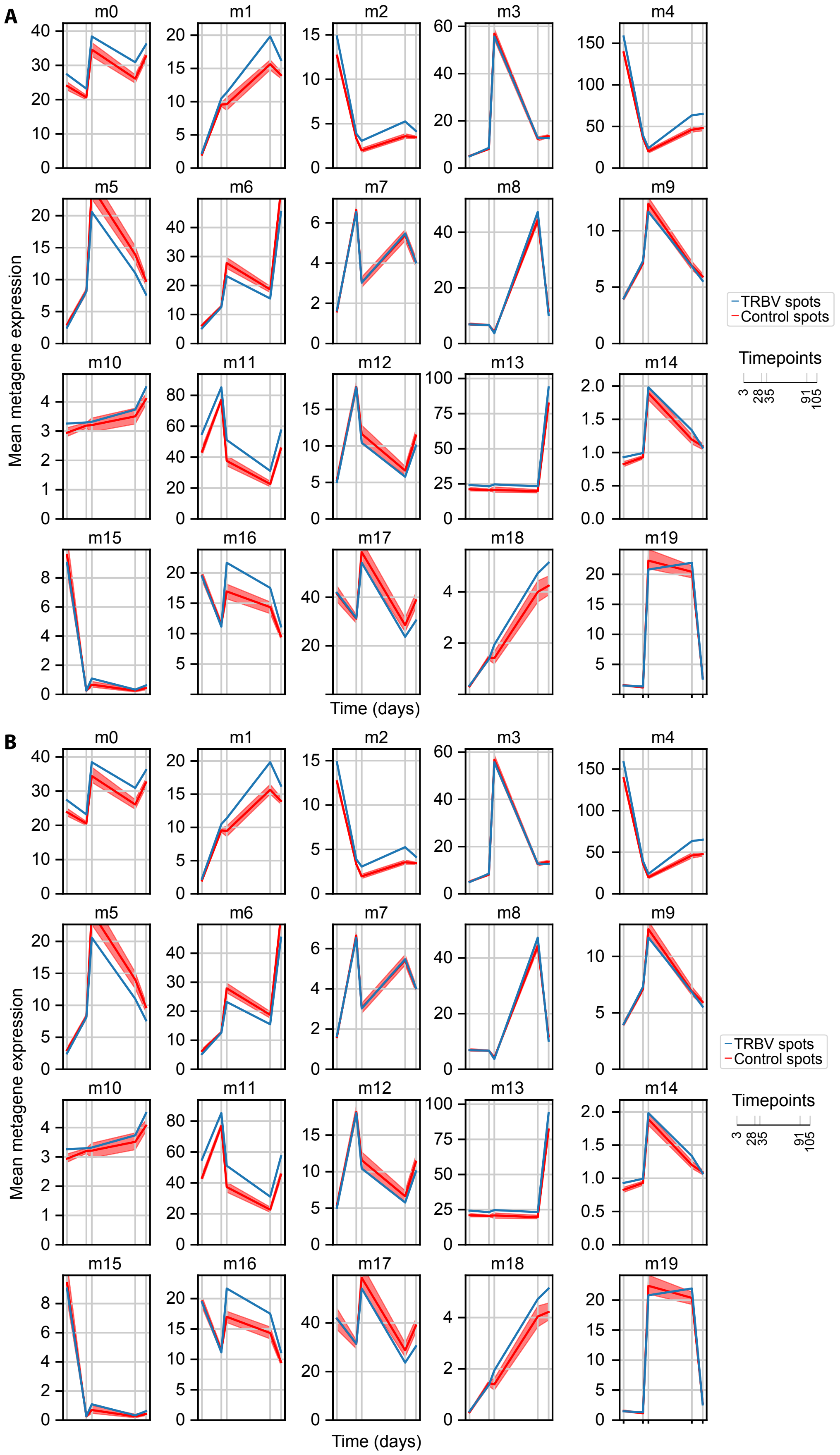

**Figure S12:** Line plot of mean embedding expression for spots with non-zero T cell receptor beta-chain variable (TRBV) usage for all metagenes in the Slide-TCR-seq data

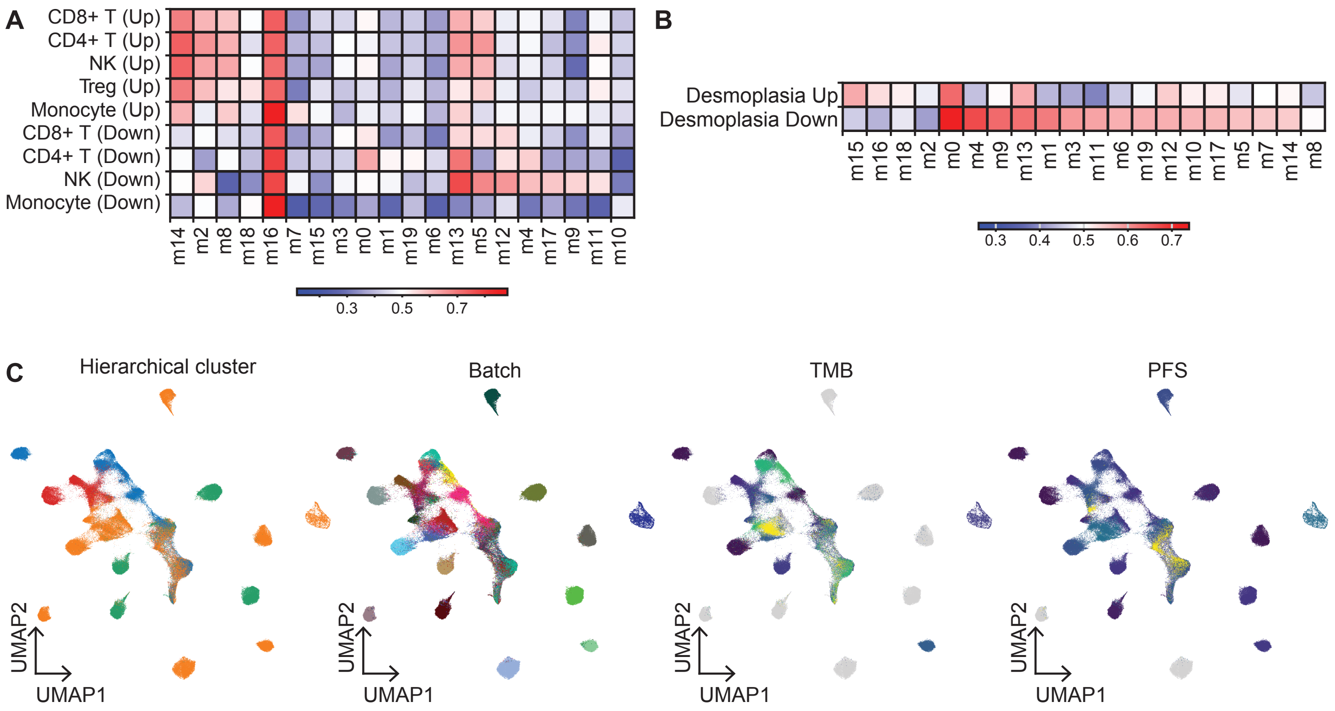

**Figure S13:** Extended analysis of metagene interpretation in the ovarian cancer dataset. **A.** AUROC scores capturing the alignment of learned Popari metagenes with predefined marker gene sets for tumor-infiltrating lymphocyte subtypes. **B.** Similar analysis as in **A.**, but for desmoplastic fibroblasts. **C.** UMAP of Popari embeddings colored by hierarchical Ward cluster (leftmost), batch (middle left), tumor mutational burden (TMB; middle right), and progression-free survival (PFS; rightmost).

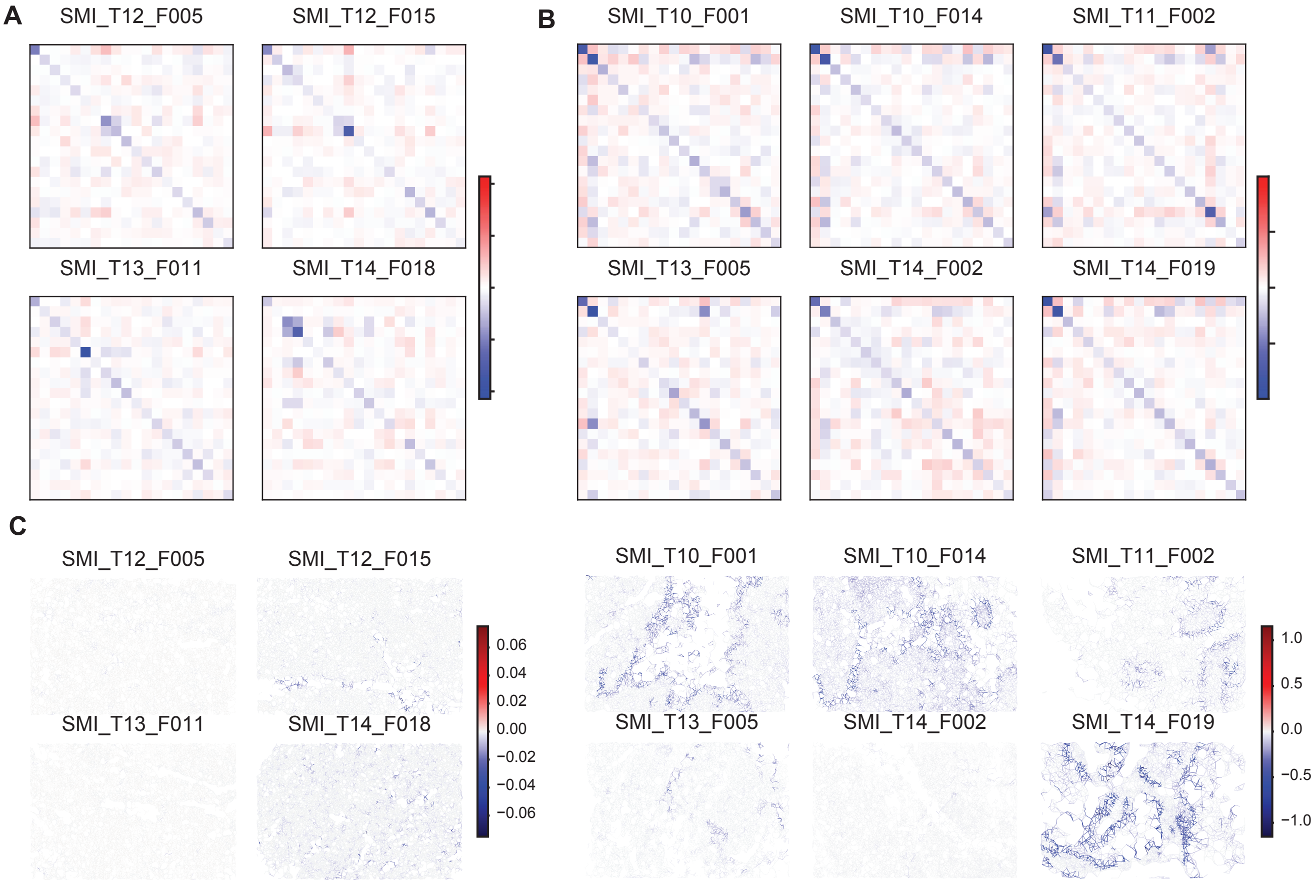

**Figure S14:** Extended analysis of distinction between malignant and stromal compartments in the ovarian cancer dataset. **A.** Spatial affinity matrices for samples in hierarchical cluster $C_4$. **B.** Similar to **A.**, but for samples in hierarchical cluster $C_2$. **C.** *In situ* $U_{x_{\text{pair}}}$ scores for metagenes m0 and m1 along edges of the spatial graph.

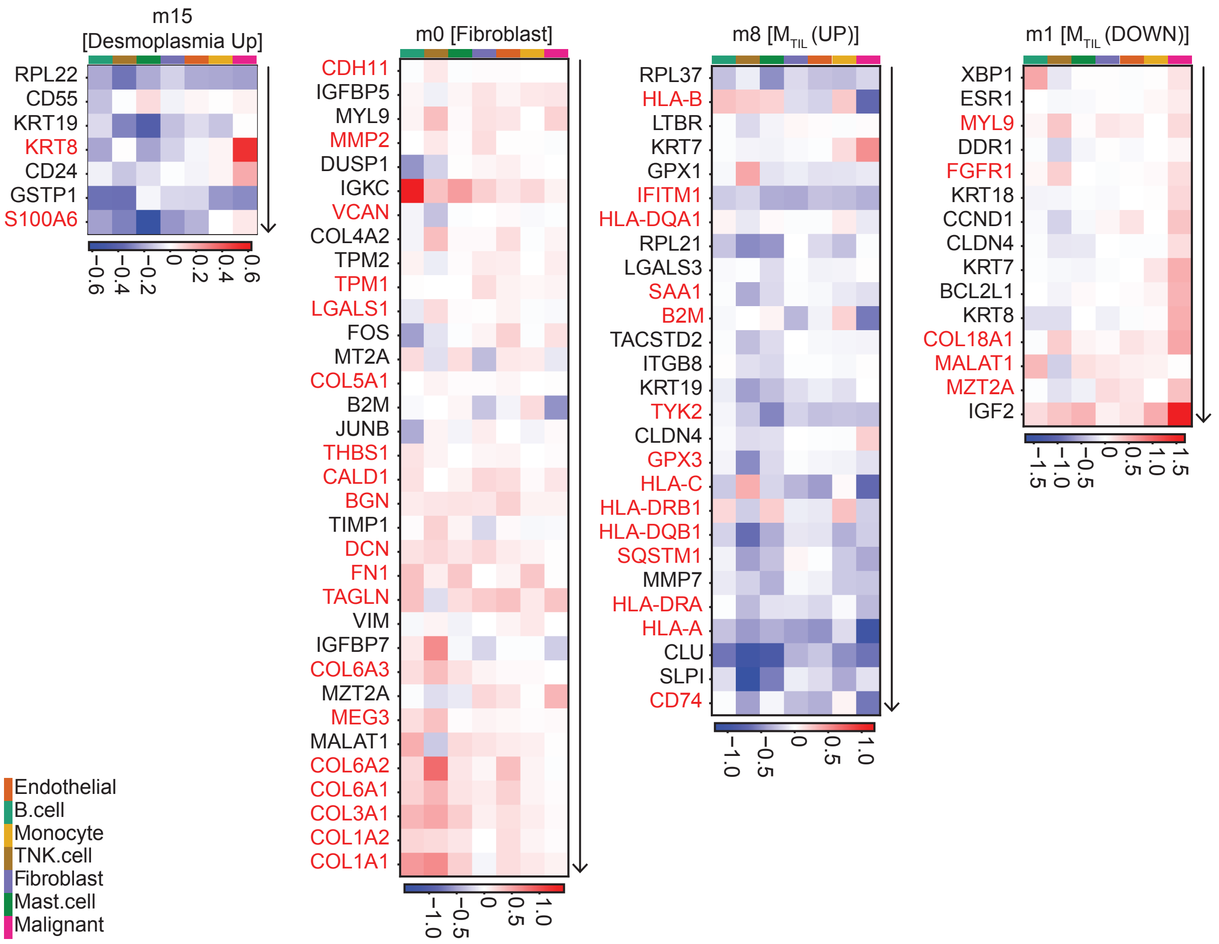

**Figure S15:** Gene expression differences between clusters $C_2$ and $C_4$ for signature genes from selected metagenes. In brackets above each signature is the gene program most strongly associated with the metagene; genes colored in red are within the gene program, and genes colored in black are not.

#### Supplementary Tables

| Broad Category | Cell Types |
|---|---|
| Antigen-presenting cells (APCs) | B cells<br>Classical type I dendritic cells (DC1)<br>Classical type II dendritic cells (DC2)<br>Monocytes (Mono)<br>Plasmacytoid dendritic cells (pDCs) |
| Effector lymphocytes (ELCs) | $CD4^{+}$ single-positive T cells (CD4+)<br>$CD8^{+}$ single-positive T cells (CD8+)<br>Intraepithelial lymphocyte precursor type A cells (IELpA)<br>Intraepithelial lymphocyte precursor type B natural killer T cells (IELpB_NKT)<br>Natural killer cells (NK)<br>Regulatory T cells (Treg)<br>Alpha-beta entry T cells (abT(entry))<br>Gamma-delta T cells (gdT) |
| Stromal cells | Endothelial cells (Endo)<br>Unknown epithelial cells (Epi_unknown)<br>Erythrocytes (Ery)<br>Fibroblasts (Fb)<br>Hematopoetic stem cells (HSCs)<br>Early thymic epithelial cells (TEC_early)<br>Vascular smooth muscle cells (VSMCs) |
| Thymocytes (TCs) | Quiescent double negative T cells (DN(Q))<br>Proliferating double negative T cells (DN(P))<br>Quiescent double positive T cells (DP(P))<br>Proliferating double positive T cells (DP(P)) |

**Table S1:** Grouping of fine-grained cell types to broad cell type categories for the Slide-TCR-seq analysis.
